# Structure-based electron-confurcation mechanism of the Ldh-EtfAB complex

**DOI:** 10.1101/2022.02.02.478877

**Authors:** Kanwal Kayastha, Alexander Katsyv, Christina Himmrichs, Sonja Welsch, Jan M. Schuller, Ulrich Ermler, Volker Müller

## Abstract

Lactate oxidation with NAD^+^ as electron acceptor is a highly endergonic reaction and some anaerobic bacteria overcome the energetic hurdle by flavin-based electron bifurcation/confurcation (FBEB/FBEC) using a lactate dehydrogenase (Ldh) in concert with the electron transferring proteins EtfA and EtfB. The electron cryo-microscopically (cryo-EM) characterized (Ldh-EtfAB)_2_ complex of *Acetobacterium woodii* at 2.43 Å resolution consists of a mobile EtfAB shuttle located between the rigid central Ldh and the peripheral EtfAB base units. The FADs of Ldh and the EtfAB shuttle contact each other thereby forming the D (dehydrogenase conducting) state. The intermediary Asp37 and Asp139 may harmonize the redox potentials between the FADs and the pyruvate/lactate pair crucial for FBEC. A plausible novel B (bifurcation conducting) state with the EtfAB base and shuttle FADs in a productive electron transfer distance was derived by integrating Alphafold2 calculations. Kinetic analysis of enzyme variants shed light on the connection between NAD binding/release and D-to-B state transition. The FBEC inactivity when truncating the ferredoxin domain of EtfA substantiates its role as redox relay. Lactate oxidation in Ldh is based on the catalytic base His423 and a metal center. On this basis, a comprehensive catalytic mechanism of the FBEC process was outlined.

## Introduction

Anaerobic fermentation processes, in particular, glucose degradation *via* glycolysis produce lactate in huge amounts (Müller, 2008). Accumulated lactate is converted to acetate by anaerobic sulfate-reducing and acetogenic bacteria (Vita et al., 2015; Weghoff et al., 2015). In the first step lactate is converted to pyruvate, the starting metabolite for further catabolic and anabolic pathways (Bock et al., 1994; Dönig & Müller, 2018; Kandler, 1983). This oxidation reaction is endergonic with NAD^+^ as electron acceptor and is driven by simultaneous co-oxidation of reduced ferredoxin (Fd_red_) by an electron-bifurcating lactate dehydrogenase/electron-transferring flavoprotein (Ldh-EtfAB) complex (Weghoff et al., 2015).

Flavin-based electron bifurcation (FBEB) is an energy-coupling process, which was developed early in evolution by anaerobic microorganisms to maximize their energy yield (Buckel & Thauer, 2013; Müller et al., 2018). In FBEB, an endergonic reaction is driven by a simultaneous exergonic redox reaction embedded into a modular, mostly soluble enzyme complex. Often, reduction of Fd_ox_ is the endergonic reaction, driven by reduction of a high potential electron acceptor. An example is the bifurcating hydrogenase (Katsyv et al., 2021; Schuchmann & Müller, 2012; Schut & Adams, 2009; Wang et al., 2013) that oxidizes hydrogen gas to protons and electrons; the latter are split and transferred to the two electron acceptors, NAD(P)^+^ and Fd_ox_. This reaction is reversible which lead to the production of H_2_ from Fd_red_ and NAD(P)H (Katsyv et al., 2021; Schuchmann & Müller, 2012; Wang et al., 2013). Thereby, the electrons flow into the reverse direction and run through a flavin-based electron confurcation (FBEC) process. The same is true for the Ldh-EtfAB complex that is electron confurcating during lactate oxidation (Weghoff et al., 2015). As key player for energy coupling in FBEB/FBEC serves a flavin endowed with an extremely inverted one-electron redox potential resulting in an extremely short-living F•^-^ state (Duan et al., 2021; Lubner et al., 2017). In the confurcation mode, the rate-determining endergonic reduction of F to F•^-^ (< - 700 mV) by Fdred (E ≈-500 mV) is strongly coupled with the exergonic reduction of F•^-^ to FH^-^ (E > +200 mV) by an electron originating from the high-potential (weak) electron donor (E < 0 mV) (Baymann et al., 2018; Nitschke & Russell, 2012). The accurate one-electron redox potentials for the Ldh-EtfAB are unknown but experimental values of E(F/F•^-^) = −911 mV and E (F•^-^/FH^-^) =+359 mV exist for the bifurcating NADPH dependent Fd:NAD^+^ oxidoreductase (Lubner et al., 2017). Strongly inverted one-electron redox potentials are also measured for quinones in the related quinone-based electron bifurcation process embedded into the *bc_1_* complex (Crofts et al., 2013). The FBEB mechanism is outlined in various comprehensive reviews (Buckel & Thauer, 2013, 2018a; Müller et al., 2018; Peters et al., 2016).

The confurcating Ldh-EtfAB complex belongs to the bEtf family of FBEB enzymes (Buckel & Thauer, 2018a), which are built up of a variable oxidoreductase/dehydrogenase core and several peripherally attached EtfAB modules (Demmer, Bertsch, Öppinger, et al., 2018; Demmer et al., 2017a; Feng et al., 2021). The EtfAB homodimer is built up of an EtfAB base composed of the N-terminal segments of EtfA and EtfB (termed domains I and III, respectively) and a module termed as EtfAB shuttle or domain II formed by the tightly associated C-terminal segment of EtfA and the C-terminal arm of EtfB (Chowdhury et al., 2014). Related structures were reported for non-bifurcating EtfAB and EtfAB-oxidoreductase complexes (Leys et al., 2003; Roberts et al., 1996; Toogood et al., 2007). Both the rigid oxidoreductase core and EtfAB base platforms and the mobile EtfAB shuttle module carry one FAD termed d-FAD, b-FAD and a-FAD. Their electrochemical potentials are unknown for Ldh-EtfAB but measured for the related Bcd-EtfAB complex except for the one-electron redox potential of the bifurcating FAD (Sucharitakul et al., 2020).

In the bEtf family a-FAD shuttles electrons between the bifurcating b-FAD and the distant d-FAD, the site of the weak electron donor (Demmer, Bertsch, Öppinger, et al., 2018; Demmer et al., 2017a) thereby creating the B (bifurcation-conducting) and D (dehydrogenase-conducting) states. The rotating EtfAB shuttle connected with the EtfAB base by two mobile linkers is holded in a defined trajectory under participation of two segments termed EtfB protrusion and EtfA hairpin (Demmer et al., 2017a). Some EtfA family members contain an N-terminal Fd domain carrying one or two [4Fe-4S] clusters (Demmer, Bertsch, Oppinger, et al., 2018) suggested to mediate ET between the external Fd and the bifurcating FAD. Deduced from the sequence and the signature motif (C(x)_17_CxxCxxC) Ldh-EtfAB possesses a Fd domain with one [4Fe-4S] cluster (Weghoff et al., 2015).

In the presented work, a comprehensive analysis of the confurcating Ldh-EtfAB reaction is performed, based on the cryo-EM structure complemented by kinetic analysis of various enzyme variants.

## Results and Discussion

### Heterologous production, purification and initial characterization of the Ldh-EtfAB complex from *A. woodii*

For structural and functional analysis the encoding strep-tagged *lctBCD* genes from *A. woodii* were cloned into the expression vector *pET21a* and expressed in the *E. coli* strain BL21(DE3)*ΔiscR* (*Supplementary File 1*). The Ldh-EtfAB complex was purified under anoxic conditions by strep-tactin affinity and Superdex 200 size exclusion chromatography. A denaturating gel showed three distinct proteins with apparent molecular masses of 51 kDa (LDH), 46 kDa (EtfA) and 29 kDa (EtfB) *(Figure 1A)* that matches to the predicted gene masses of 51.2 kDa, 46.2 kDa and 29.1 kDa. Native gel electrophoresis and analytical gel filtration data revealed a rather fragile multimeric complex with maximal molecular masses of 220 kDa *(Figure 1B)* and 280 kDa, respectively. Gelfiltration profiles clearly indicate the partial dissociation of the Ldh-EtfAB complex into Ldh and EtfAB.

**Figure 1.**
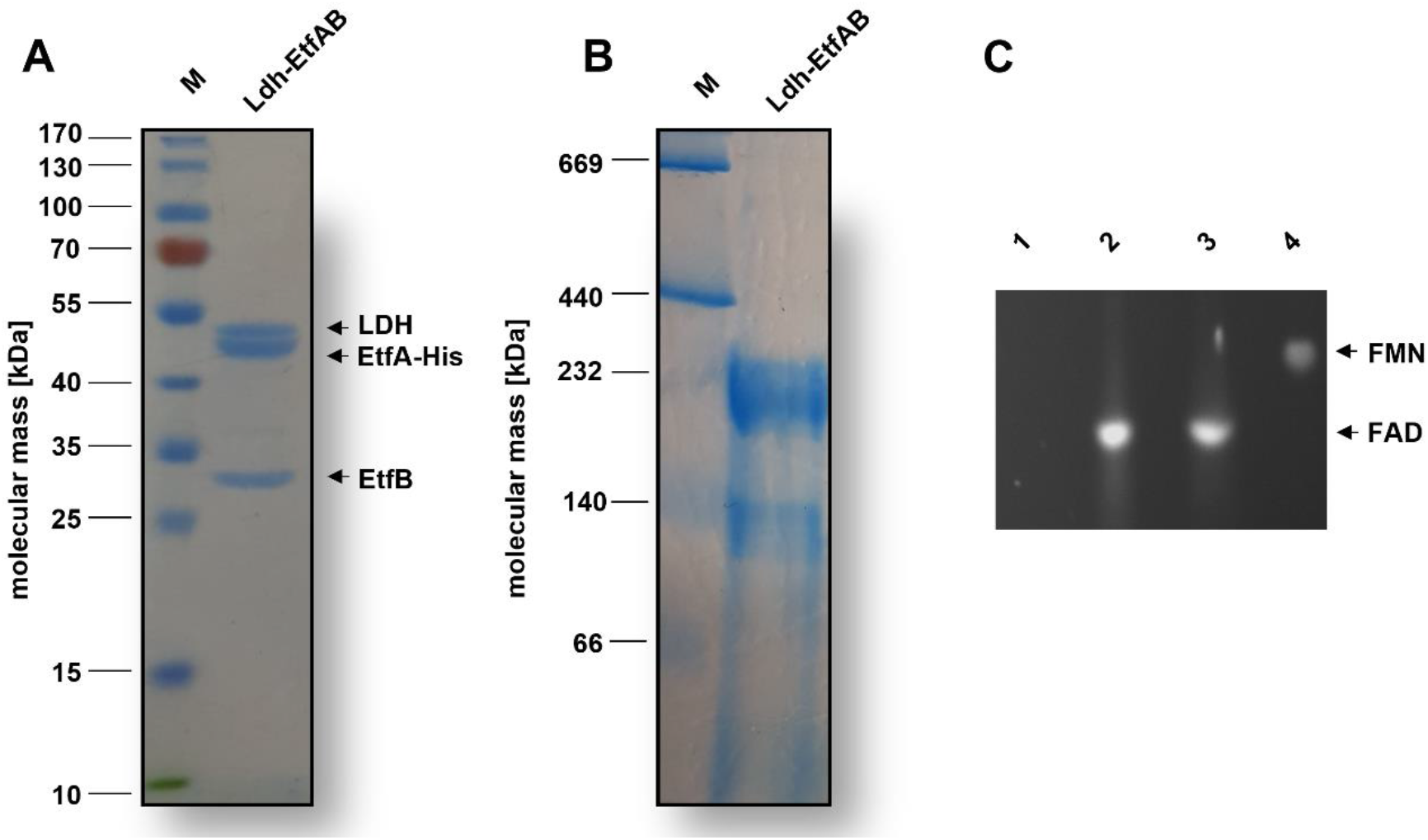
SDS-, native PAGE and flavin determination of the purified Ldh-EtfAB complex. The Ldh-EtfAB complex was anaerobically purified by strep-tactin affinity and Superdex 200 size exclusion chromatography. 5 μg or 20 μg of the purified protein was separated by a 12% SDS-(**A**) or native (**B**) PAGE. Coomassie Brilliant Blue G250 was used for protein staining. (**C**) Flavins of ~ 1 nmol Ldh-EtfAB were separated on a TLC plate using 60% [v/v] n-butanol, 15% [v/v] glacial acetic acid and 25% [v/v] H_2_O as the mobile phase. 1 nmol of FAD and FMN was used as standards. For this experiment Ldh-EtfAB was purified with buffers additionally mixed with 5 μM FAD and FMN. lane 1, buffer; lane 2, FMN; lane 3, FAD; lane 4, Ldh-EtfAB; M, Prestained PageRuler™ **Source data 1.**Source data for Figure 1

The heterologously produced Ldh-EtfAB complex shows similar biochemical properties as the protein isolated directly from *A. woodii* (Weghoff et al., 2015). It catalyzes the confurcating Fdred-dependent lactate:NAD^+^ oxidoreductase reaction with a rate of 12.3 ± 1.8 U/mg. and the bifurcating pyruvate-dependent NADH:Fd_ox_ oxidoreductase reaction with a rate of 1.2 ± 0.2 U/mg. Confurcation activity was optimal between 25 and 30°C *(Figure 2A)*. It decreased below 25 or above 30°C by 40 or 20% and at 45°C to minor values. The complex had his highest confurcation activity at pH 7 *(Figure 2B)*. A decrease of the FBEC activity to 50% could be observed at pH 6 and 8 and inactivity at pH 5 and 10. Similar FBEC activities could be measured, when separately produced Ldh (data not shown) and EtfAB (separated in the gelfiltration column) were added together, which demonstrate the stability of the individual components and their capability to assemble.

**Figure 2.**
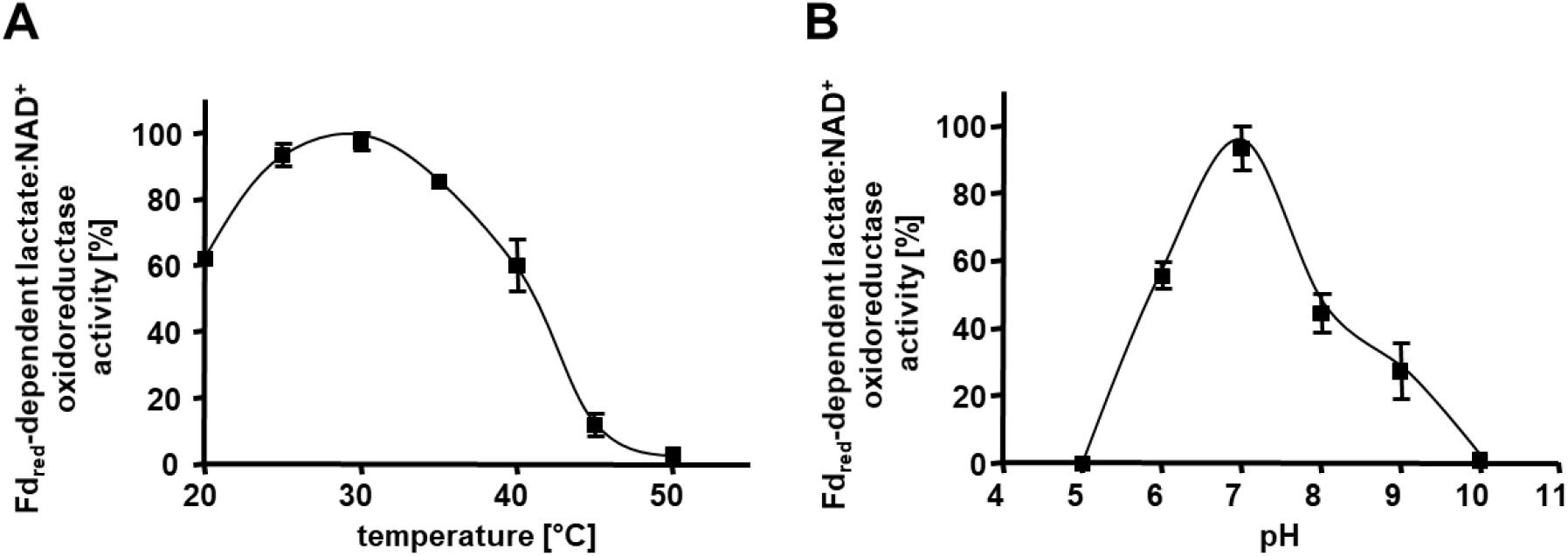
Basic biochemical properties of the Ldh-EtfAB complex. The temperature-(**A**) and pH-(**B**) optima for the Ldh-EtfAB complex were determined. The assay contained 30 μM Fd, 17 μg CODH, 4 mM NAD^+^ and 5 μg LDH-EtfAB in a 100% CO gas atmosphere. The buffer used for temperature optima was 50 mM Bis-Tris, 50 mM CaCl_2_, pH 7.0. After 5 min incubation at the appropriate temperature, the reaction was started by adding 50 mM D/L-lactate. Buffers used for the pH optima determination contained 50 mM MES, 50 mM CHES, 50 mM CAPS, 50 mM Bis-Tris, 50 mM Tris and 50 mM CaCl2. The reduction of NAD^+^ was measured by absorption spectroscopy at 340 nm. The data represents the mean and standard deviation of two independent experiments (n = 2), each performed in triplicate.

### Global structural features of the Ldh-EtfAB complex

The cryo-EM structure of the fragile Ldh-EtfAB complex from *A. woodii* was determined from a sample cross-linked with BS^3^, because multiple attempts without cross-linking failed. From 9788 micrographs and 674283 extracted particles collected a twofold-averaged density map at a mean resolution of 2.43 Å was calculated *(Figure 3A, Figure supplement 1A and B)* using Relion (Scheres, 2012). The local resolution ranges from 2.3 Å for the Ldh core to 3.1 Å for a peripheral EtfA region *(Figure 3B)*. Only for the N-terminal Fd domain (2-62) of EtfA the EM map was highly disordered and model building was impossible. The polypeptide chain of all other structural parts could be largely traced with ARP/WARP (Langer et al., 2008). Missing residues were afterwards manually incorporated with COOT (Emsley & Cowtan, 2004).

**Figure 3.**
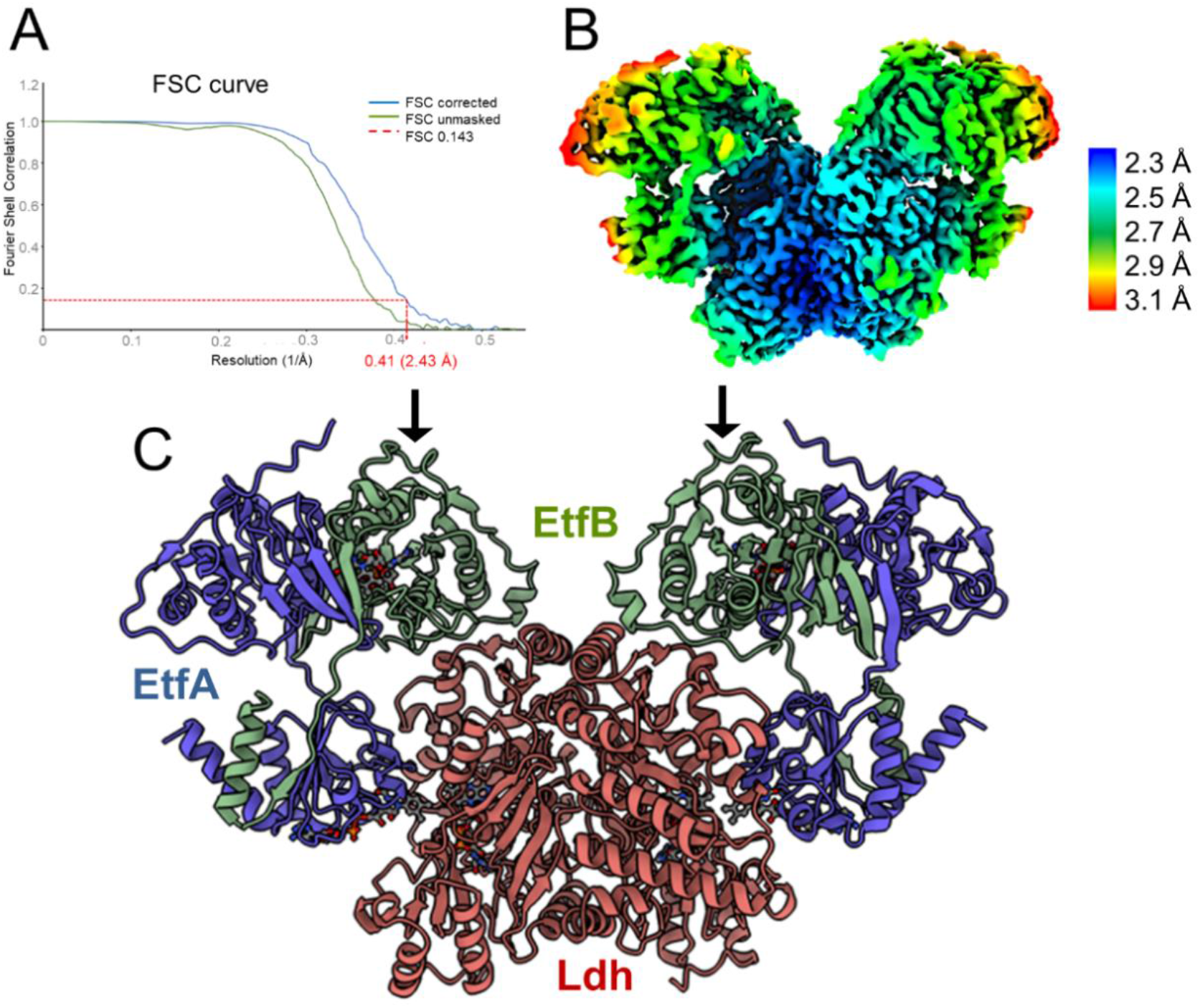
Structure of (Ldh-EtfAB)2 complex in the D-state. (**A**) Fourier Shell Correlation curve of unmasked and corrected 3D maps at 0.143 “gold standard” (**B**) Cryo-EM map at 2.43 Å resolution. The density is colored according to the local resolutions. (**C**) Ribbon representation. The Ldh dimer is drawn in red, EtfA in blue and EtfB in green. The 250 kDa heterohexameric complex has an overall size of 150 Å x 100 Å x 60 Å. The docking sites of the two Fd-like domain are marked with black arrows. **Figure – Supplement 1A.** EM data (A) Negative-staining. (B) Selected 2D classes foo 3D map reconstruction. (C) Particle distribution on a C2-symmetry EM map. (D) Relion image processing working flow. **Figure – Supplement 1B.** Table for EM statistics.

The Ldh-EtfAB complex was found as a heterohexamer composed of a Ldh dimer forming the core and two EtfAB modules peripherally associated. Exclusively, the Ldh subunits provide the interface between the two Ldh-EtfAB protomers *(Figure 3C)*. The active site regions of the two Ldh-EtfAB protomers are ca. 30 Å apart from each other and most likely operate independently. The derived molecular mass of 252 kDa complex agrees with the value derived from gelfiltration and native PAGE analysis. In accordance with the X-ray structures of the Bcd-EtfAB/CarCDE complexes (Demmer, Bertsch, Öppinger, et al., 2018; Demmer et al., 2017a) the cryo-EM (Ldh-EtfAB)2 structure shows both Ldh-EtfAB protomers in the resting D (dehydrogenase conducting) state *(Figure 4A)* in which electrons are exchangeable between a-FAD and d-FAD. During the reaction cycle electron transfer (ET) has also to be adjusted between a-FAD and b-FAD implicating a rotation of the EtfAB shuttle towards the B (bifurcation conducting) state *(Figure 4B)*. Analysis of the EM micrographs with the aim to find a small population in the B state failed. Accordingly, bifurcating oxidoreductase-EtfAB complexes are, so far, only experimentally trapped in the D state therefore considered as stable resting conformation while the B state appears to to be short-living and transient. In contrast, non-bifurcating oxidoreductase-EtfAB complexes contain a promiscuous EtfAB and form a transient interface between the EtfAB shuttle and their dehydrogenase partners which is beneficial for sampling a wider conformational space (Leys et al., 2003).

**Figure 4.**
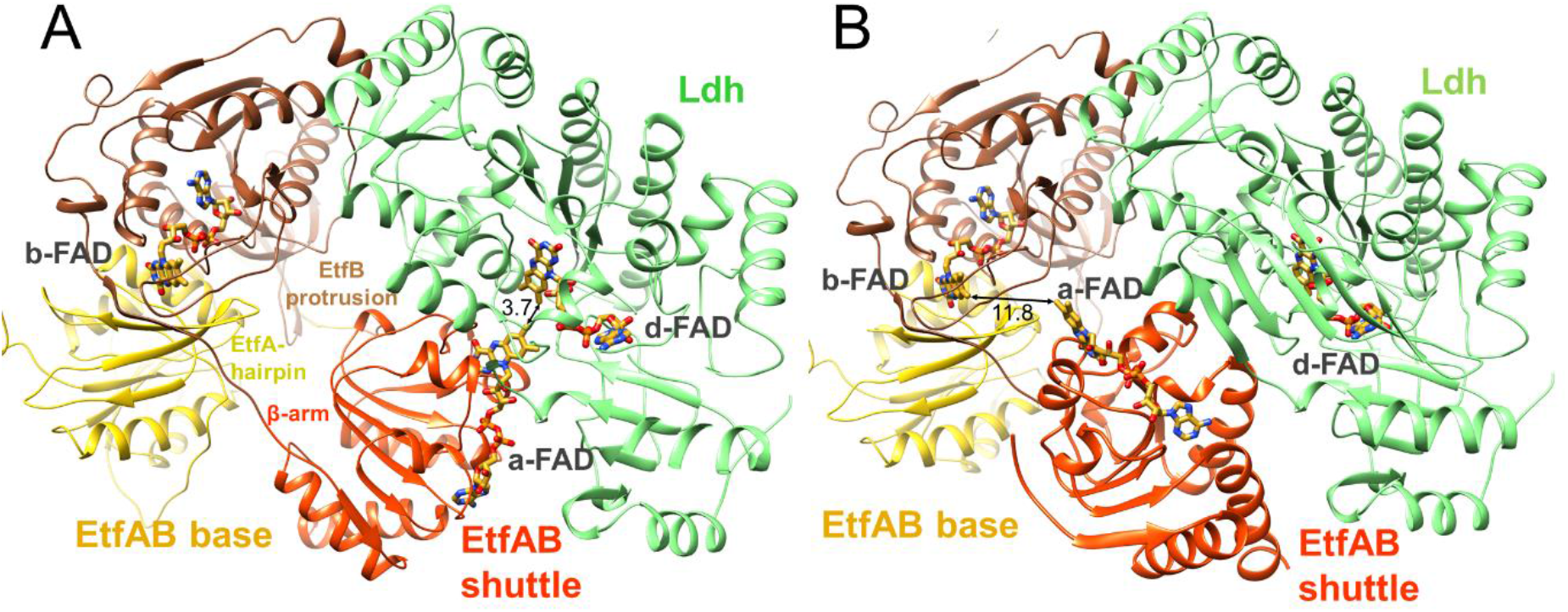
Structural states of the (Ldh-EtfAB)2 complex. (**A**) The experimentally characterized D-state. The EtfAB shuttle (red) is spatially separated from the rigid EtfAB base (EtfB domain III in yellow and EtfA domain I in brown) and attached to Ldh (green). a-FAD and d-FAD are 3.7 Å, a-FAD and b-FAD ca. 37 Å apart from each other. (**B**) The B-state. By combining the experimental EM data and Alphafold2 calculations a solid B-state could be established for the first time. Starting from the D-state the EtfAB shuttle swings 75° to adjust an ET distance between a-FAD and b-FAD. The shortest distance between the isoalloxazine rings is 11.8 Å. Upon rotation the contact to Ldh is lost while that to the EtfAB base is formed. **Figure – Supplement 1.** The Ldh-EtfAB interface. (A) Contact area between Ldh and the EtfAB base. (B) Contact region between Ldh and domain II in the D-state. **Figure – Movie 1**. Transition between the D- and B-states

The oscillation of the EtfAB shuttle between the D- and B-states is realized by a fixed interface between the Ldh core and the EtfAB base and a variable interface formed either between the EtfAB shuttle and Ldh or between the EtfAB shuttle and the EtfAB base *(Figure 4)*. The fixed interface is constituted by the small segment 185-193 of EtfB composed of an elongated loop and a short helix which is strictly conserved in bifurcating and non-bifurcating EtfBs and referred to as recognition loop *(Figure 4A – Figure supplement 1)* (Toogood et al., 2007). The variable interface in the D-state is centered around a-FAD and d-FAD. In the Ldh-EtfAB complex the nonpolar xylene moiety of the isoalloxazine rings point towards each other; the edge-to-edge distance is 3.7 Å *(Figure 4A – Figure supplement 1)*. For adjusting a productive interflavin distance the variable contact region of the D state has to be individually adapted for different EtfAB dehydrogenases /oxidoreductases. A special situation is reported for the Fix/EtfABCX complex as ET between the EtfAB shuttle and FixC is mediated by the additional Fd subunit FixX (Feng et al., 2021). The B-state could be, so far, not experimentally trapped but a model of the EtfAB structure integrating the recently published Alphafold2 (Jumper et al., 2021) offered a convincing result *(Figure 4B)*. Surprisingly, the EtfAB shuttle does not move to the conducting site starting from the isolated EtfAB structure (Chowdhury et al., 2014; Demmer et al., 2017b) by further rotation but flows on a different trajectory by a 75°rotation starting from the D-state *(Figure 4 – Movie 1)*. a-FAD becomes attached to the elongated loop linking β-strands 201:206 and 213:217 of EtfB and the EtfB protrusion (*Figure 4B*). The xylene rings of b-FAD and a-FAD point to each other and the shortest distance between their methyl groups is 11.8 Å.

### The Ldh subunit and its active site

Ldh is built up of two domains, an N-terminal FAD domain (1-218) subdivided into two α + β subdomains and a cap domain (219-417) characterized by an antiparallel β-sheet element as well as an extended C-terminal arm (418-467) attached to the FAD domain (*Figure 5A*). This fold classifies Ldh as a member of a flavoenzyme family with p-cresol methyl hydroxylase (CMH) (Cunane et al., 2000), vanillyl-alcohol oxidase (Mattevi et al., 1997), MurB (Benson et al., 1997) and membrane-associated Ldh (Dym et al., 2000) as prototypes (Fraaije et al., 1998); their corresponding rms deviations from Ldh of *A. woodii* are 2.6 Å (1DIQ, 407 of 467 residues used), 2.6 Å (1AHU, 399 of 467), 3.5 Å (2MBR, 203 of 467) and 2.4 Å (1FOX, 371 of 467). Except for the membrane-associated Ldh CMH-type flavoenzymes are homodimers. D-FAD is embedded between the two α + β subdomains except for the isoalloxazine ring that protrudes beyond the FAD domain towards the cap domain. Therefore, the isoalloxazine is well accessible from bulk solvent (*Figure 5A)*.

**Figure 5.**
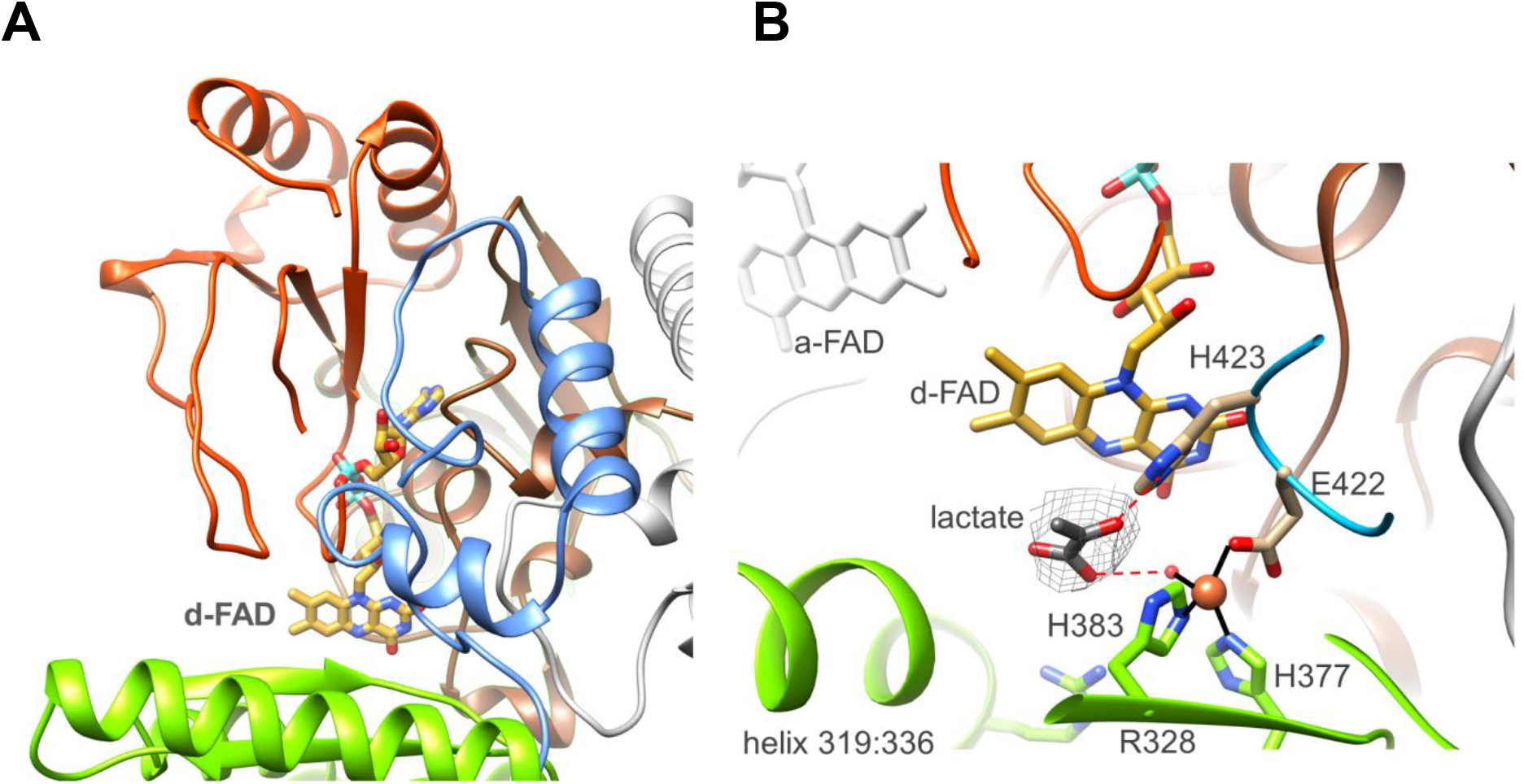
Ldh. (**A**) The overall structure. The Ldh subunit belongs to the CMH flavoenzyme family composed of two α + β subdomains (orange-red and brown), a cap domain (lightgreen) and a C-terminal extension (blue). d-FAD (as stick with carbon in yellow) is positioned between them. The isoalloxazine ring is fixed by several van der Waals contacts and two hydrogen-bonds between FAD N3C2O and Gly153CONH and between FAD O4 and Gly138NH of Ldh. (**B**) The active site. Lactate was tentatively modelled into a density (gray) in front of the N5 of d-FAD involving a metal site for its binding. Its hydroxy group appears to be activated by His423. We tentatively assigned the metal as Fe^2+^ (orange) due to its addition during purification. **Figure – Supplement 1.** Sequence alignment of Ldh

In front of its *si*-side the substrate binding site can be easily reached by lactate/pyruvate *via* a wide gate framed by Gly138, Met155, Gly329 and Leu332 (*Figure 5B)*. The substrate binding sites is essentially formed by Glu44, Asp346, His377, Asn382, His384, Tyr386, Glu422 and His423 directly or indirectly involved in lactate binding/ oxidation and well conserved in the membrane-associated Ldh (*Figure 5B – Figure supplement 1*) (Dym et al., 2000). Notably, we detected a putative metal binding site in the substrate binding cavity ligated by His377, His384, Glu422 and a water molecule. An unexplained density in front of d-FAD corresponds to the profile of pyruvate or lactate (*Figure 5B)*. Lactate was modelled in a manner that the hydride transferring carbon is 3.6 Å apart from N5 and the hydroxy group is hydrogen-bonded with the invariant His423 (*Figure 5B – Figure supplement 1*) perhaps acting as a catalytic base during lactate oxidation. The carboxyl group of the substrate would be fixed by interactions with the metal-ligating water and Arg328.

### The EtfAB module carrying b-FAD

The bifurcating EtfAB module of the (Ldh-EtfAB)2 complex is structurally related to other family members reflected in rms deviations between its EtfAB base (domain I, 62-269 and domain III, 2-236) and those of the Bcd-EtfAB, CarCDE and Fix-EtfABX of ca. 1.4 Å (5ol2 (Demmer et al., 2017a), 404 of 443), 1.4 Å (6FAH (Demmer, Bertsch, Öppinger, et al., 2018), 416 of 443) and 2.4 Å (7KOE (Feng et al., 2021), 368 of 443). The corresponding values for the EtfAB shuttle (EtfA 270-398, EtfB: 237-265) are ca. 1.4 Å (155 of 158), 1.8 Å (153 of 158) and 2.1 Å (112 of 158).

b-FAD is considered as the heart of FBEC/FBEB and exchanges electrons with a-FAD, Fd and NAD (Chowdhury et al., 2014). For evaluating its binding and catalytic function crucial residues were substituted by the single nucleotide exchange method *via* corresponding primers (*Supplementary File 1*). For all enzyme variants presented the yield and subunit composition correspond to that of the wild type enzyme (*Table 1 – Table supplement 1A)*. The oxidoreductases-EtfAB including Ldh-EtfAB contain the conserved ^122^DGDTAQVGP^130^ stretch as recognition motif for b-FAD binding (*Table 1 – Table supplement 1B*), which is absent in non-bifurcating Etfs (Buckel & Thauer, 2018b; Chowdhury et al., 2014; Garcia Costas et al., 2017). The [Δb-FAD] variant consisting of the D122A, D124A, T125G, Q127G, V128A and P130A substitutions shows a lower overall FAD amount of 1.7 ± 0.4 mol per mol enzyme. The FBEC and FBEB monitoring Fd_red_-dependent lactate:NAD^+^ and pyruvatedependent NADH:Fd_ox_ oxidoreductase activities were reduced to 7% and not anymore measurable, respectively. Likewise, the NADH:DCPIP oxidoreductase activity, measuring the hydride transfer from NADH to b-FAD, was also abolished. Both findings can be explained by the expected strong decrease of the b-FAD content. On the other hand, lactate to pyruvate oxidation with ferricyanide as electron acceptor is only reduced by a factor of 2 in the [Δb-FAD] variant (*Table 1*) indicating two independently acting active sites. Moreover, we exchanged the conserved residue R205 of EtfA (*Table 1* – *Table supplement 1B)* that forms a hydrogen-bond to N5 of b-FAD. As found for the equivalent CarCDE variant (Demmer, Bertsch, Öppinger, et al., 2018) the [ΔR205] variant of Ldh-EtfAB completely losts the confurcating/bifurcating capabilities (*Table 1*). Although all analyzed FBEB enzymes share a positively charged arginine/lysine residue (Chowdhury et al., 2014; Demmer et al., 2015; Wagner et al., 2017; Watanabe et al., 2021) in contact with N5 of b-FAD its specific function beyond FAD binding is still obscure (Kayastha et al., 2021). However, the maintained NADH:DCPIP oxidoreductase activity exclude larger rearrangements of the FAD and the adjacent polypeptide and thus substantiate the crucial importance of the N5 environment for the unusual redox behaviour of the flavin. In the B-state D189 is 11 Å apart from b-FAD (*Table 1 – Table supplement 6)* as well as 4.5 and 10 Å apart from the old a-FAD position and the new one calculated by Alphafold2 (Jumper et al., 2021). The moderate catalytic change of the ΔD189 variant expressed in a 35% and 15% decrease of the confurcating and bifurcating activity rather argues for a FAD binding site farer from D189 and thus also for the new B-state. As a control [ΔR205] and [ΔD189] variants were still capable to reduce ferricyanide or DCPIP with lactate or NADH (*Table 1*).

**Table 1.** Specific activities of recombinant Ldh-EtfAB and site-specifically changed variants with artificial electron donors/acceptors. **Table – Supplement 1A.** SDS-PAGE of the site-specifically changed Ldh-EtfAB variants.
**Table – Supplement 1B.** Location of the EtfA and EtfB mutations. **Source data 1**. Source data for Table 1 -Supplement 1A

|  | WT | ΔFe/S | ΔFe/S-arm | ΔD189 | ΔR205 | Δb-FAD | ΔNAD<sup>+</sup> | ΔSPT |
| --- | --- | --- | --- | --- | --- | --- | --- | --- |
| **Fd<sub>red</sub>:lactate: NAD<sup>+</sup> (340 nm)** | 12.3 ± 2.3 | 0.1 ± 0.05 | 0.1 ± 0.06 | 7.7 ± 2.6 | 0.01 ± 0.01 | 0.9 ± 0.3 | 0.2 ± 0.1 | 1.1 ± 0.3 |
| **NADH:pyruvate:Fd<sub>ox</sub> (340 nm)** | 1.2 ± 0.2 | 0.5 ± 0.03 | 0.1 ± 0.1 | 0.9 ± 0.1 | 0.02 ± 0.02 | 0.06 ± 0.02 | 0.01 ± 0.01 | 0.01 ± 0.05 |
| **lactate:K<sub>3</sub>[Fe(CN)<sub>6</sub>] (420 nm)** | 19.0 ± 5.2 | 13.9 ± 5.4 | 14.6 ± 4.8 | 11.1 ± 6.7 | 15.8 ± 4.3 | 8.3 ± 9.2 | 8.5 ± 7.3 | 8.7 ± 6.5 |
| **NADH:DCPIP (600 nm)** | 8.8 ± 2.5 | 7.3 ± 0.2 | 9.7 ± 1.5 | 5.6 ± 2.5 | 6.3 ± 2.3 | 0.4 ± 0.2 | 0.2 ± 0.1 | 9.2 ± 0.4 |

\*NAD<sup>+</sup>, Fd, ferricyanide or DCPIP reduction was measured at 340, 430, 450 or 600 nm, respectively. Values are given as specific activities in U/mg protein.

To substantiate the assumed binding site for NAD^+^/NADH (Chowdhury et al., 2014) we prepared the [ΔNAD] variant by substituting R87, F89 and G91 to alanine (*Table 1 – Table supplement 1B)*. NADH oxidation in the confurcation activity assay was reduced to 8 % of the wildtype and the NAD^+^ reduction in the bifurcation activity assay was no longer measurable.

In addition, DCPIP could not be reduced by NADH anymore but ferricyanide with lactate with a two-fold lower activity (8.5 ± 7.3 U/mg) (*Table 1)*. Altogether, the vital role of b-FAD as bifurcating flavin and of NADH as its hydride donor/acceptor was demonstrated independent from structural interpretation.

Finally, we exchanged the highly conserved residues S223, P224 and T225 localized at the anchor of the EtfB arm in the vicinity of the NAD binding site (*Table 1* – Table supplement 1B). This [ΔSPT] variant showed 90 % lower Fd_red_-dependent lactate:NAD^+^ and almost completely lost pyruvate-dependent NADH:Fd_ox_ oxidoreductase activities (*Table 1)*.The lactate:K_3_Fe(CN)_6_ and DCPIP: NADH oxidoreductase activities were 50 % lower than those of the native enzyme indicating that mutations primarily do not affect the individual redox reactions but their coupling. In the context of previous data (Demmer, Bertsch, Öppinger, et al., 2018; Demmer et al., 2017a; Schut et al., 2019a) these amino acid exchanges disturb the transmission of NAD^+^ binding towards the EtfB arm and thus impair EtfAB shuttle rotation.

### The Fd domain

EtfA of the Ldh-EtfAB complex contains a Fd domain with one [4Fe-4S] cluster (Weghoff et al., 2015) in contrast to the structurally characterized EtfA subunits of Bcd-EtfAB and CarCDE endowed with no N-terminal Fd or an N-terminal Fd domain with two [4Fe-4S] clusters, respectively (*Figure 6A*) (Chowdhury et al., 2014; Demmer, Bertsch, Oppinger, et al., 2018). In the EM structure the density of the Fd domain in both Ldh-EtfAB protomers is highly disordered such that chain tracing and also the detection of the [4Fe-4S] cluster is not feasible. Its location could, however, be identified close to the position determined by Alphafold2 (Jumper et al., 2021). Accordingly, the preserved [4Fe-4S] cluster in the Ldh-EtfAB complex is the one closer to b-FAD when compared with the superimposed CarCDE. Its distance of 18 Å to b-FAD is still a bit too long for ET (*Figure 6B*). For functional analysis the entire Fd domain (2-65) was truncated or cysteines 41, 44 and 47 coordinating the [4Fe-4S] clusters were exchanged to alanine. In these [ΔFe/S-domain] and [ΔFe/S] variants the irons are completely lost (0.3 ± 0.2 and 0.5 ± 0.3 mol Fe /mol protein); the stability and subunit composition were, however, not affected (*Table 1 – Table supplement 1B*) and the overall yield increases up to four-fold compared to the native Ldh-EtfAB complex. Kinetic analysis revealed that the FBEC/FBEB activities were abolished in both variants (*Table 1*) whereas the functionality of the isolated active sites is maintained. The DCPIP:NADH oxidoreductase activity of the [ΔFe/S] or [ΔFe/S-arm] variants was 7.3 ± 0.2 and 9.7 ± 1.5 U/mg, respectively, and the lactate:K_3_Fe(CN)_3_ oxidoreductase activity 13.9 ± 5.4 and 14.6 ± 4.8 U/mg, respectively (*Table 1*), both values being 25% lower than those of the wildtype enzyme.

**Figure 6.**
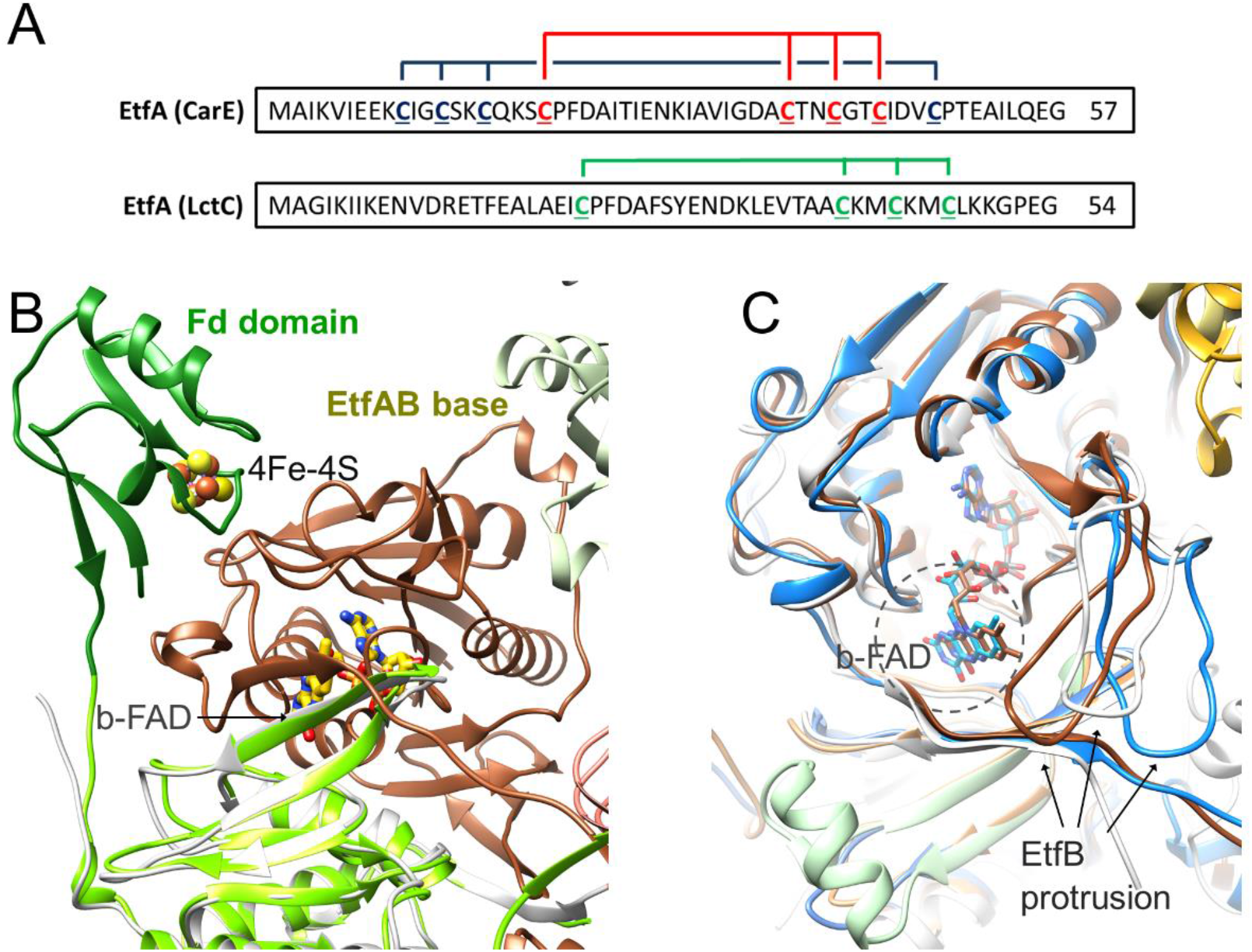
Ferredoxin (Fd). **(A**) The sequence of the N-terminal Fd domain of the CarCDE and the Ldh-EtfAB complexes. EtfA of the Ldh-EtfAB complex contains one [4Fe-4S] cluster ligated with four cysteines marked in green. In comparison, the equivalent CarE of the CarCDE complex possesses two [4Fe-4S] clusters marked in blue and red. The latter is shared between CarCDE and Ldh-EtfAB. (**B**) The structure of the Ldh-EtfAB complex superimposed with EtfA (green) including the Fd domain (dark green) calculated by Alphafold2. The Fd domain interacts with EtfB via the segment linking the anchor of the EtfB arm (225-232) and the terminal strand (203:207) of the central β-sheet and via the loop following strand 81:85. The [4Fe-4S] cluster (in ball-and-stick) of the Fd domain was modelled based on the properly positioned four cysteine sulfurs. (**C**) Accessibility of b-FAD. b-FAD is more shielded in Ldh-EtfAB (red) than in the superimposed CarCDE (blue) due to the reoriented EtfB protrusion such that an arriving Fd might not find a docking site sufficiently close to b-FAD. The structure of non-bifurcating EtfAB of *C. propionicum* (grey) (calculated by Alphafold2) was also superimposed to visualize the position of the inserted segment (lightgreen). An interference with an external Fd remains elusive. The approximate diameter of Fd is drawn as dashed circle.

In contrast, the CarCDE complex preserves the FBEB capability after cutting off its Fd domain. We, therefore, conclude that external Fd can be placed next to b-FAD in truncated CarCDE but not in the [ΔFe/S] or [ΔFe/S-arm] variants of Ldh-EtfAB. Superimposed Ldh-EtfAB and CarCDE reveals a displacement of the EtfB protrusion by nearly 15 Å, which might impair Fd binding close to b-FAD in Ldh-EtfAB (*Figure 6C*). In summary, these result shows the vital importance of the [4Fe-4S] custer in Ldh-EtfAB as specific adaptor for external Fd binding. It is worth mentioning that the non-bifurcating EtfABs from *Clostridium propionicum*(Cprop2325) highly related to their bifurcating partners contain an insertion region that may block Fd, as already predicted (Buckel & Thauer, 2018b), in a manner related to Ldh-EtfAB (*Figure 6C*).

### Enzymatic mechanism

Biochemical and kinetic studies on the recombinant Ldh-EtfAB complex indicated a 10 times higher enzymatic activity for the FBEC than for the FBEB process. Therefore, the proposed mechanism is outlined in this direction (*Figure 7*) starting from the D-state. (1) Lactate binds into a pocket in front of the isoalloxazine of d-FAD and is oxidized to pyruvate *via* a hydride transfer to N5 forming d-FADH^-^. The active site is formed by residues conserved in FBEC and membrane-spanning Ldhs (Dym et al., 2000) (*Figure 5 – Figure supplement 1*) suggesting a related enzymatic mechanism (2). One electron of d-FADH^-^ rapidly flows to a-FAD thereby generating d-FADH• (d-FAD•^-^) and a-FAD•^-^ (a-FADH•). It is worth to mention that a-FAD might be already present as flavosemiquinone in the resting state of the cell and would be reduced in this case to a-FADH^-^. (3) NAD^+^ binds into a pocket at the *si*-side of b-FAD as verified by site-directed mutagenesis experiments (*Table 1*) in agreement with various previous studies (Chowdhury et al., 2014). In the characterized D-state Thr226 and Val228 of EtfB occupy the frontside of b-FAD and have to be pushed aside upon NAD^+^ binding (*Figure 6C*). According to reports on Bcd-EtfAB, CarCDE and Fix-EtfABCX complexes (Chowdhury et al., 2014; Demmer, Bertsch, Oppinger, et al., 2018; Schut et al., 2019b) and supported by mutational analysis (see above) NAD^+^ binding substantially influences the conformation of the EtfB anchor and thus may influence the adjustment of the B and D states. (4) External Fd binds to the mobile Fd domain of EtfA in a manner to enable rapid ET between their two [4Fe-4S] clusters. As b-FAD is with 18.4 Å a bit too far away from the [4Fe-4S] cluster of the Fd domain (according to the EtfA structure calculated by Alphafold2), external Fd_red_ binding might induce conformational changes to reduce their distance to ca. 14 Å. (5) Perhaps induced by NAD^+^ and/or Fd_red_ binding the EtfAB shuttle rotates into the B-state (*Figure 4 – Movie 1*). (6) Fd_red_ fires one electron *via* the [4Fe-4S] cluster of EtfA to the bifurcating b-FAD (*Figures 6A and 7A*). (7) The produced energy-rich b-FAD•^-^ is instantaneously reduced to b-FADH^-^by a-FAD•^-^ of the EtfAB shuttle arrested in the B state (*Figure 4B*). According to general electrochemical principles of FBEB both one-electron donation processes to b-FAD accompanied by protonation are strictly coupled (*Figure 7B*) (Baymann et al., 2018; Nitschke & Russell, 2012). (8) The nicotinamide ring of NAD^+^ attached parallel to the isoalloxazine ring becomes reduced by transferring a hydride from N5 of b-FADH^-^ to C4 of NAD^+^. (9) Upon release of NADH and/or Fd_ox_ the EtfAB shuttle swings back into the D-state and a-FAD uptakes the remaining electron of d-FADH• (d-FAD•^-^). In analogy to the first round b-FAD is reduced in the B-state by one-electron transfers from Fd_red_ and a-FAD•^-^ rotated again from the D-into the B-state (*Figure 4 – Movie 1*). Finally, a second NAD^+^ is reduced to NADH (*Figure 7A*).

**Figure 7.**
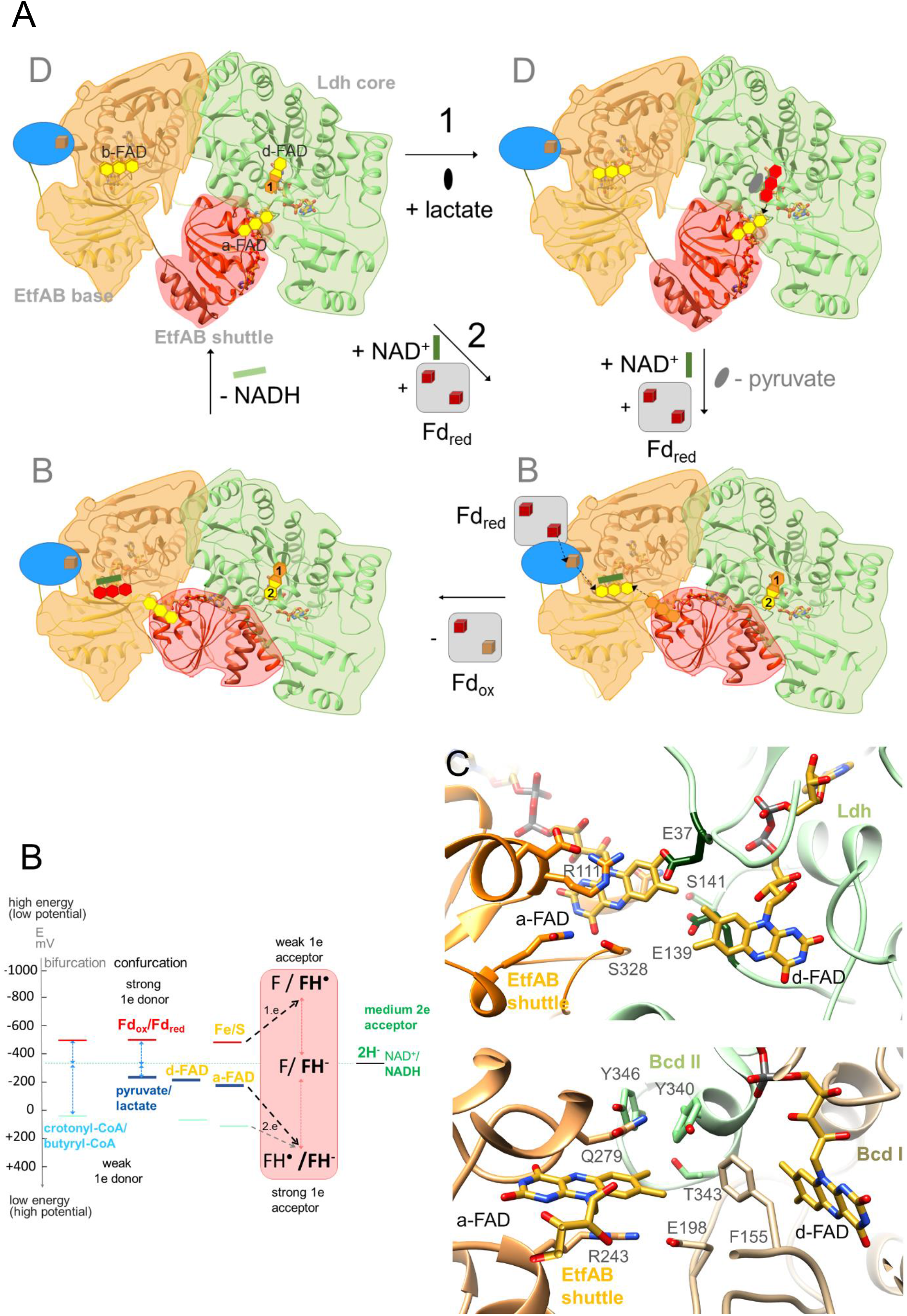
The Ldh-EtfAB reaction. (**A**) The mechanism. The reaction cycle was outlined in the direction of the thermodynamically preferred confurcation direction. It starts from the structurally characterized D state and runs through two rounds (termed 1 + 2). The three-membered rings of FAD, FAD•^-^ / FADH• and FADH^-^ are drawn in yellow, red and orange. The d-FAD isoalloxazine is shown with orange and yellow moieties representing in the 1. round FADH• (FAD•^-^) and in the 2. round FAD. The Fd domain is drawn as blue ellipse and the external Fd as grey box. (**B**) Thermodynamic scheme of FBEC. Low-potential Fd_red_ donates an electron to b-FAD in a slow endergonic reaction that is driven by the exergonic ET from a-FAD•^-^ to b-FAD•^-^. This escapement-type mechanism may prevent short circuit reactions e.g. from b-FAD•^-^ to a-FAD•^-^ and undesirable side reaction of the highly reactive flavosemiquinone (Baymann et al., 2018). The generated b-FADH^-^ transfers a hydride to NAD^+^. A confurcation reaction occurs when the redox potential difference from the medium electron acceptor (NAD^+^) to the weak electron donor (lactate) is smaller than to the strong electron donor (Fd_red_). (**C**) Microenvironments of a-FAD and d-FAD in the D-state of Ldh-EtfAB (upper panel) and Bcd-EtfAB (lower panel). In Ldh-EtfAB the two isoalloxazine rings are closer and the protein surrounding is significantly different compared to Bcd-EtfAB/CarCDE. Asp37 and Asp139 contacting a-FAD and d-FAD may destabilize an electron-rich reduced state resulting in a lower redox potential (*Figure 7B*).

The Ldh-EtfAB complex catalyzes the first dominantly electron-confurcating reaction structurally studied. As the overall direction of the process is thermodynamically driven, the reaction cycle becomes just reversed. The midpoint redox potential of the pyruvate/lactate pair is −190 mV, which is substantially lower than that of the crotonyl-CoA/butyryl-CoA pair of - 10 mV, which turns a bifurcation into a confurcation event (*Figure 7B*). When assuming that the redox potentials of the electron carriers between b-FAD and lactate also differ by ca. 180 mV the microenvironment of a-FAD and d-FAD should be distinguished between Bcd-EtfAB and Ldh-EtfAB (*Figure 7C*). The most striking features represent the acidic residues Glu37 and Glu139 adjacent to the isoalloxazines of a-FAD and d-FAD that are only found in the Ldh-EtfAB complex and may destablize the electron-rich reduced state. In addition, the surrounding of a-FAD of Ldh-EtfAB compared to Bcd-EtfAB reveals by a histidine-to-phenylalanine exchange at position 348 despite the high overall conservation among bEtfB family members. Thus, the redox potentials of both FADs might therefore be decreased towards the value of the pyruvate/lactate pair (*Figure 7B and C*).

## Materials and methods

### Cloning of lctBCD and generation of site-specifically mutated variants

The genes *lctBCD* (3507 bp) were amplified from chromosomal DNA using the primers lctBCD_pET21a_for and lctBCD_pET21a_rev (*Supplementary File* 1B) and cloned into the expression vector *pET21a* (5406 bp), amplified using the primers pet21a_for and pet21a_rev, *via* Gibson Assembly (New England Biolabs) (*Supplementary File 1A*). A sequence encoding a Strep-tag was introduced at the 3’-end of the gene *lctC* coding for EtfA by using corresponding primers (*Supplementary File 1B*). The resulting plasmid *pET21α_lctBC-StrepD* was the template for site directed mutagenesis. Nucleic acid changes were introduced using corresponding primers at desired loci in the encoding genes (*Supplementary File 1B*). Generated plasmids were checked by sequencing. *pET21a* plasmids were transformed into *Escherichia coli* BL21(DE3) *ΔiscR*.

### Production and purification of the Ldh-EtfAB complex in E. coli BL21(DE3)ΔiscR

*E. coli* BL21(DE3)Δ*iscR* was grown aerobically in modified LB-medium (1% Trypton, 1% NaCl, 0,5% yeast extract, 100 mM MOPS, pH 7.4) supplemented with 4 mM ammonium-iron(II)citrate and 25 mM glucose at 37°C. At an OD_600_ of 0.5 – 0.8 the culture was transferred into a sterile, anaerobic Müller-Krempel flask (Glasgerätebau Ochs, Bovenden-Lenglern, Germany) and supplemented with 2 mM cysteine and 20 mM fumarate. After closing the culture with a butyl stopper, the flask was further incubated at 16 °C. As soon as the culture was cooled down to 16°C gene expression was induced by addition of IPTG to a final concentration of 1 mM. All steps from this point onward were executed in an anaerobic chamber (Coy Laboratory Products, Grass Lake, USA) with a mixed gas phase of N_2_/H_2_ (95:5 [v/v]). After 16 h - 19 h cells were anaerobically harvested, resuspended in 200 mL buffer W (50 mM Tris, 150 mM NaCl, 20 mM MgSO_4_, 20 % (v/v) glycerol, 4 μM resazurin, 5 μM FAD, 2 mM DTE, pH 8) and disrupted once in a French press (SLM Aminco, SLM Instruments, USA) after addition of DNase I and 0.5 mM PMSF (setting high, 1000 psi). The lysate was centrifuged (14000 x g, 20 min, RT) to separate undisrupted cells from the crude extract. Subsequently, the Ldh-EtfAB complex was purified by affinity chromatography on Strep-Tactin high-capacity material (IBA Lifesciences GmbH, Göttingen, Germany) using the elution buffer E (buffer W+ 5 mM desthiobiotin). Fractions containing Ldh-EtfAB were pooled and concentrated *via* ultrafiltration (Vivaspin 6, 30 kDa Cut-off, Sartorius Stedim Biotech GmbH, Göttingen, Germany) to a volume of 500 μl. The concentrate was separated with a flow rate of 0.5 ml/min on a Superdex 200 10/300 GL column (GE Healthcare, Little

Chalfont, UK), previously equilibrated with buffer W. For cryo-EM studies buffer S (20 mM HEPES, 150 mM NaCl, pH 7.5) was used for the size exclusion step. Afterwards, the Ldh-EtfAB complex was stabilized with bis(sulfosuccinimidyl)suberate (BS^3^) crosslinker (Thermo Fisher Scientific, Waltham, USA) by incubating the sample (≈ 1 mg) with 1 mM BS^3^ for 20 min at room temperature. The assay was stopped by addition of 50 mM Tris. To get rid of impurities, the crosslinked-sample was again separated with a flow rate of 0.5 ml/min on a Superdex 200 10/300 GL column (GE Healthcare, Little Chalfont, UK), previously equilibrated with buffer S.

### Assays of lactate dehydrogenase activity

Kinetic measurements were routinely performed in a N_2_ atmosphere at 30 °C in 1.8ml anaerobic cuvettes (Glasgerätebau Ochs), which were sealed with rubber stoppers. Physiological Fd_red_-dependent lactate:NAD^+^ - and pyruvate-dependent NADH:Fd_ox_ oxidoreductase activities are measured as previously described (Weghoff et al., 2015). The same assay conditions were used for recording the lactate:K3Fe(CN)6- and NADH:DCPIP oxidoreductase activities using 1 mM K_3_Fe(CN)_6_ and 500 μM DCPIP (dichlorophenol indophenol), respectively. The NAD^+^/NADH (ε = 6.3 mM^−1^ cm^−1^), Fd (purified from *C. pasteurianum* (Schönheit et al., 1978)) (ε = 13.1 mM^-1^ cm^-1^), DCPIP (ε = 20,7 mM^-1^ cm^-1^) or K_3_Fe(CN)_6_ (ε = 1 mM^-1^ cm^-1^) oxidation/reduction was monitored at 340, 430, 600 or 420 nm by UV/Vis spectroscopy, respectively.

### Analytical methods

Protein concentration was measured according to Bradford (Bradford, 1976) The protein complex was separated by PAGE, using the SDS buffer system of Laemmli (Laemmli, 1970) or Wittig (Wittig et al., 2007) and stained with Coomassie brilliant blue G250. The molecular mass of the purified Ldh-EtfAB complex was determined using a calibrated superdex 200 column, buffer E and defined size standards (ovalbumin: 43 kDa; albumin: 158 kDa; catalase: 232 kDa; ferritin: 440 kDa). The iron content of the purified enzyme was determined by colorimetric methods (Fish, 1988) and the flavin content by thin layer chromatography (TLC) as described before (Bertsch et al., 2013).

### Single-particle electron cryo-microscopy

For sample vitrification C-Flat R1.2/1.3 copper, 300-mesh grids (Electron Microscopy Sciences) were glow discharged thrice with a PELCO easiGlow device at 15 mA for 45 seconds. 4 μl of protein with a concentration of 1.8 mg/ml were applied to a freshly glow-charged grid, blotted for seconds at 4°C, 100 % relative humidity and a blot force of +20. Grids were vitrified by plunging into liquid ethane using a Vitrobot Mark IV device (Thermo Scientific).

Movies were recorded using a Titan Krios G3i microscope operated at 300 kV (Thermo Scientific) and equipped with a Gatan BioQuantum imaging filter using a 30 eV slit width and K3 direct electron detector. Data were collected at a nominal magnification of 105,000 × in electron counting mode using aberration-free image-shift (AFIS) correction in EPU (Thermo Scientific). Applied parameters are listed in Figure 4 – Supplement 1B.

After using CryoSPARC live (Punjani et al., 2017) for on-the-fly processing data to check data quality the full dataset was processed using RELION-3.1 (Scheres, 2012; Zivanov et al., 2018). Beam-induced motion was corrected using MOTIONCOR2 (Zheng et al., 2017) and dose-weighted images were generated from movies for initial image processing. Initial CTF parameters for each movie were estimated using Gctf algorithms (Zhang, 2016). Particles were picked with crYOLO (Wagner et al., 2019) using the neural network trained general model approach and cleaned *via* 2D classification in RELION-3.1. Unsupervised Initial model building, 3D classification, 3D refinement, along with post-processing steps like CTF refinement, Bayesian polishing and final map reconstructions were performed with RELION-3.1. Maps were visualized with Chimera (Pettersen et al., 2004) and models automatically built with Arp/Warp (Langer et al., 2008) or manually within COOT (Emsley & Cowtan, 2004). Further refinement was performed using PHENIX (Adams et al., 2010) and the quality of the model was assessed with COOT and MolProbity (Williams et al., 2018). Figures 3 B+C, 4, 5, 6 and 7C were produced with Chimera.

## Acknowledgments

Work in the laboratory of V.M. was supported by the Deutsche Forschungsgemeinschaft (DFG). K.K. thanks the International Max Planck Research School and Hartmut Michel for funding. We further thank Hartmut Michel, Janet Vonck and Werner Kühlbrandt for generous support.

## Additonal information

### Author contributions

V.M. and U.E. designed the research. The enzyme was purified by A.K., and the EM structure determined by K.K. C.H. and A.K. cloned the *lctBCD* variants. A.K. heterologously expressed the enzyme and did the mutational analysis. S.W. provides the cryo-EM facility and technical support. J.M S. was involved in cross-linking. K.K., A.K., U.E. and V.M. wrote the paper.

### Additional files

Figure 3 - Supplement 1

Figure 4 -Supplement 1

Figure 5 -Supplement 1

Table 1 – Supplement 1

Supplementary File 1

Raw data. Movie 1

Source data 1 for Figure 1

Souce data 1 for Table 1 – Supplement 1A

Source data 1 for Supplementary File 1

### Data availability

The data supporting the finding of this study are available from the corresponding authors upon reasonable request. The cryo-EM map and coordinates were deposited under accession numbers EMD-13960 and 7QH2, respectively.

---

**Figure 3 – Figure supplement 1.**
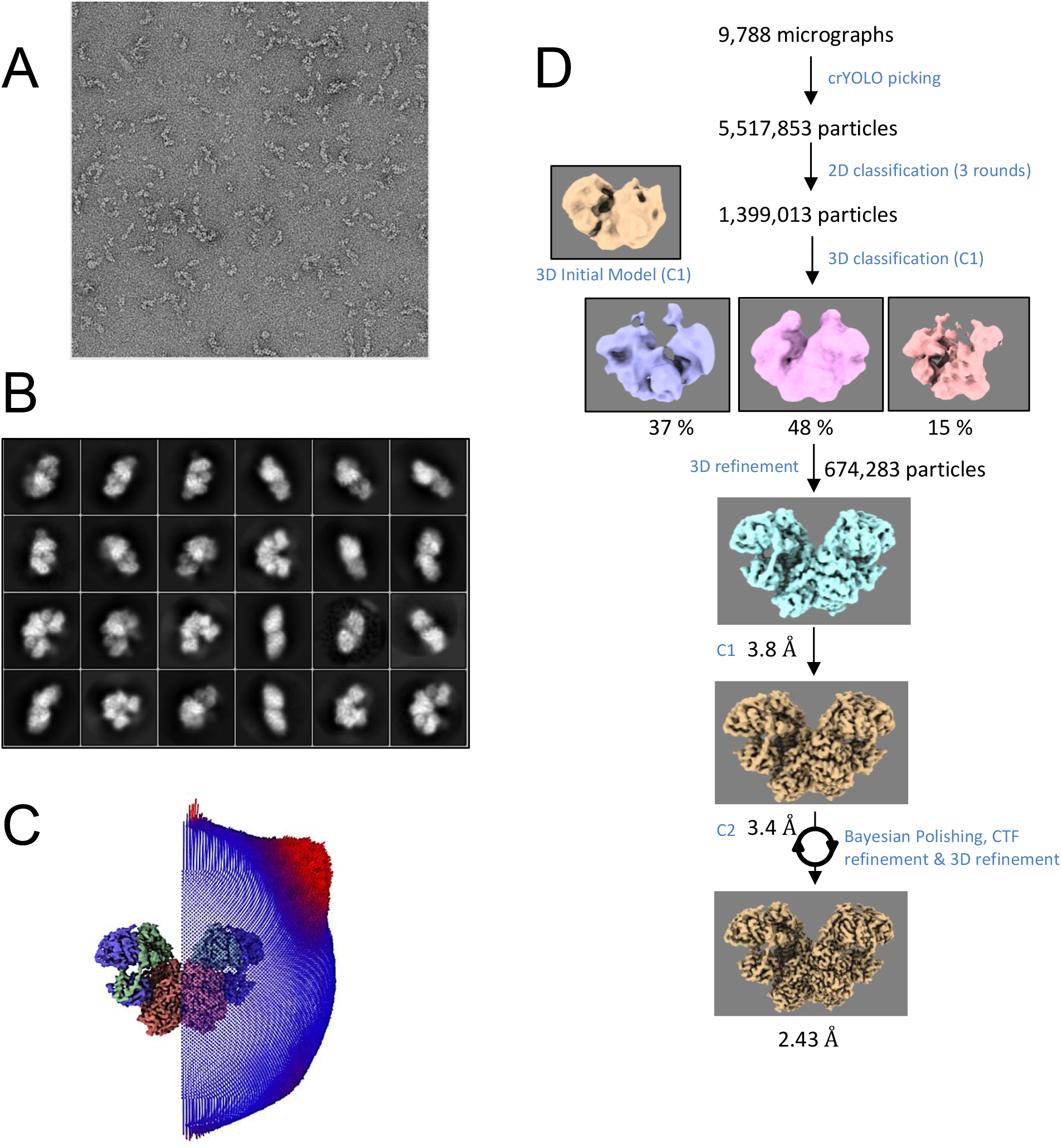
EM data. (A) Negative-staining Electron Microscopy. For screening the quality of the sample, negative-staining was performed with ca. 82 nM protein and stained using 0.75 % uranyl formate for about 90 seconds. Grids used were QuantiFoil R1.2/1.3 Cu-200-mesh coated with collodion solution (cellulose nitrate, 2% in amyl acetate, Sigma) and then sputter-coated with 4.33 nm carbon thread for the final carbon film for protein sample support and distribution. Before applying the sample, grids were glow-discharged twice with a PELCO easiGlow device at 15 mA for 45 seconds. (B) Selected 2D classes for 3D map reconstruction. (C) Particle distribution on a C2-symmetry EM map. (D) Relion image processing work flow used for the structure determination of the Ldh-EtfAB complex.

**Figure 3 – Supplement table 1.** EM statistics

| **Experimental data** |  |
| --- | --- |
| Protein | Ldh-EtfAB complex |
| State | Resting D-state |
| **Data collection and Processing** |  |
| Microscope | Thermo Scientific Titan Krios G3i |
| Voltage (kV) | 300 |
| Camera | Gatan K3 summit |
| Exposure time (s) | 6.52 |
| Total dose (e<sup>-</sup>/Å<sup>2</sup>) | 106.17 |
| Dose per frame (e<sup>-</sup>/Å<sup>2</sup>) | 1.01 |
| Defocus range (μm) | 1.2-2.1 |
| Pixel size (Å) (calibrated) | 0.837 |
| Magnification (nominal) | 105,000x |
| Symmetry imposed | C2 |
| No. of micrographs | 9,788 |
| Initial particle number | 5,517,853 |
| Final particle number | 674,283 |
| Map resolution (Å) | 2.43 |
| FSC threshold | 0.143 |
| **Refinement** |  |
| Map-sharpening B factor (Å<sup>2</sup>) | -67 |
| Model composition |  |
| Chains | 6 |
| Protein (Residues) | 2,140 |
| Ligands | FAD: 6; FE: 2 |
| RMSD Bond Length (Å) | 0.005 |
| RMSD Bond Angles (°) | 1.17 |
| MolProbity score | 2.77 |
| Clash score | 4.39 |
| Ramachandran plot (%) |  |
| Favored | 97.0 |
| Allowed | 2.9 |
| Outliers | 0.1 |
| Rotamer outliers (%) | 1.9 |
| ADP (B-factor) (min/max/mean) |  |
| Protein | 55.1/151.3/90.6 |
| Ligand | 60.6/93.4/79.2 |

**Figure 4 – Figure supplement 1.**
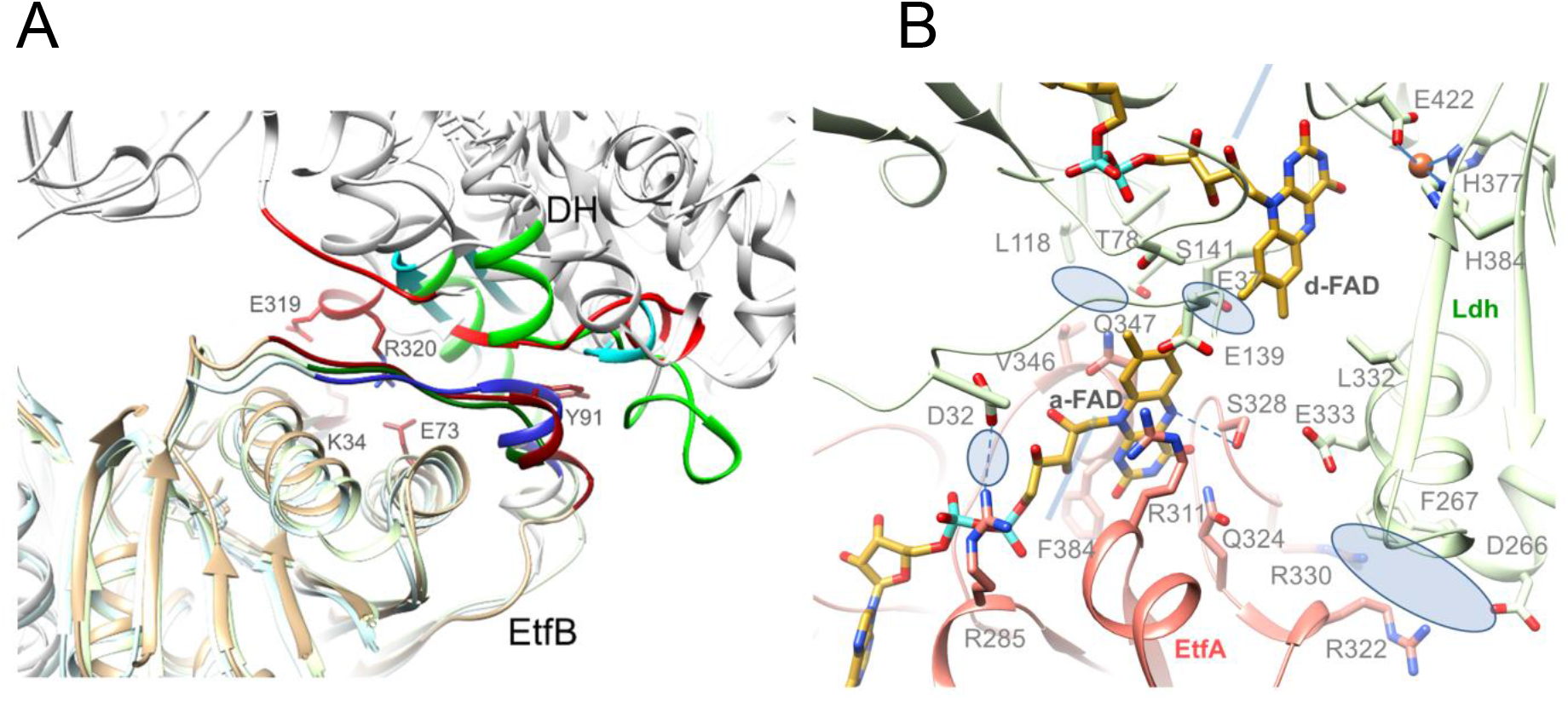
Ldh-EtfAB interface. (**A**) Contact area between Ldh and the EtfAB base. As reported for non-bifurcating EtfAB the side chain of Tyr119 of EtfB protrudes into the hydrophobic pocket formed by Leu195 and Tyr297 of EtfB, and Ala301, Val312, Ile314 of Ldh. In addition, a second small hydrophobic patch (Leu188 of EtfB and Pro273, Leu277, Tyr297, Val315 and Val324 of Ldh) and two remote salt bridges (Lys34 of EtfB - Glu319 of Ldh, Glu73 of EtfB - Arg320 of Ldh) are formed. (**B**) Contact region between Ldh and domain II in the D-state. The major contact point is formed around the a-FAD and d-FAD, which are in van der Waals contact to each other. Above and underneath the central flavin interface lies three rather small hydrophobic and hydrophilic contact patches (marked by blue ellipses), respectively, involving the FAD and cap domains of Ldh and a single salt bridge between Arg285 of EtfA and Asp32 of Ldh.

**Figure 5 – Figure supplement 1.**
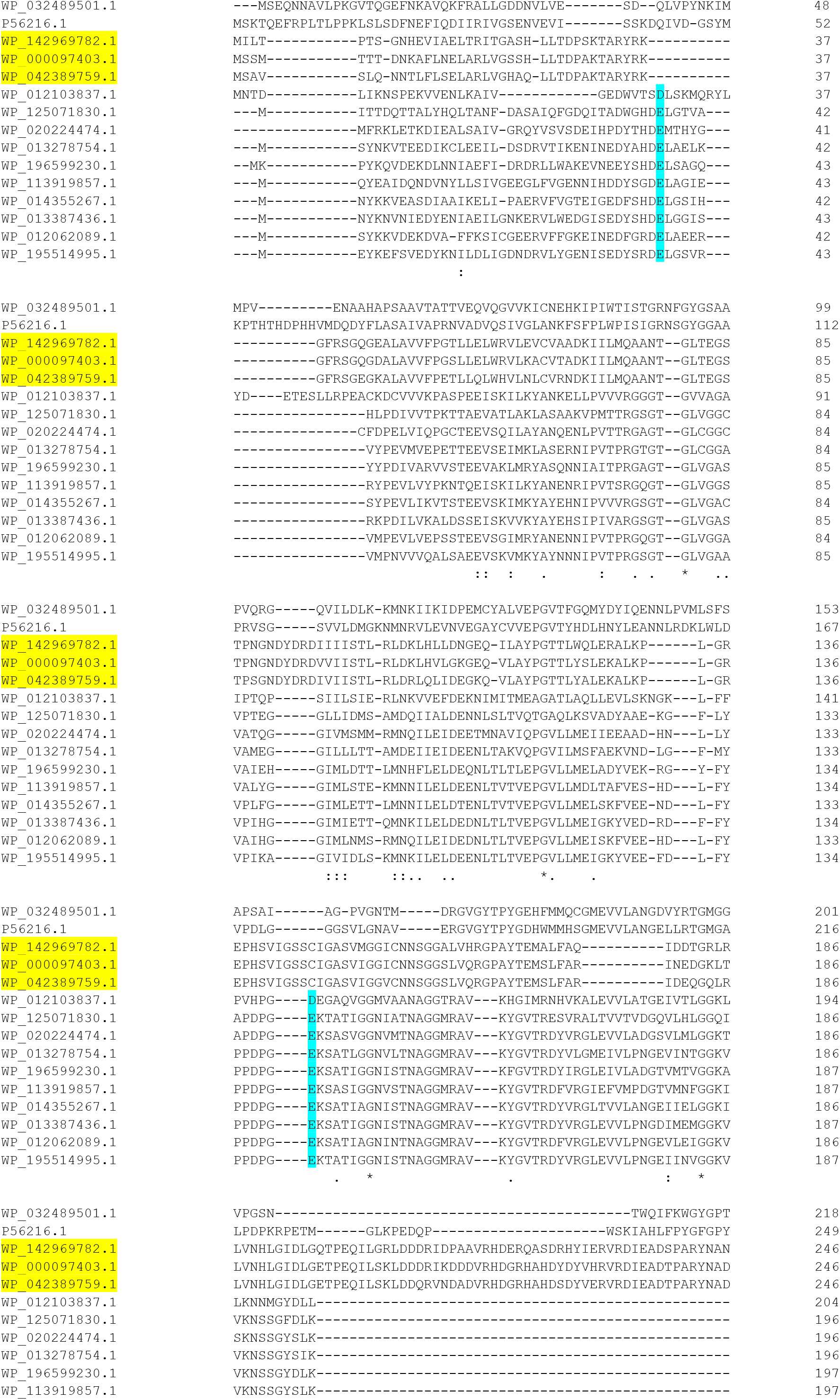
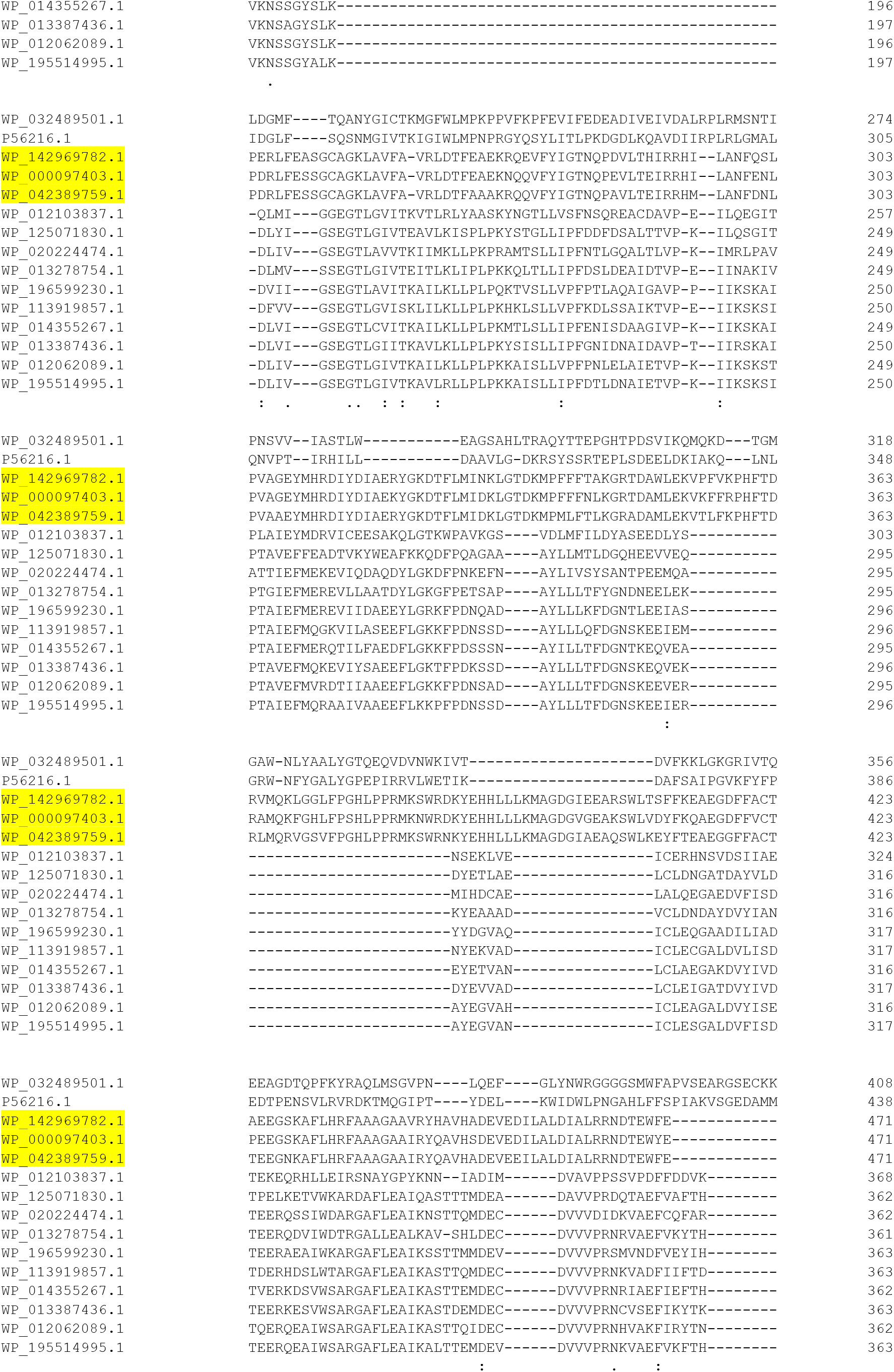
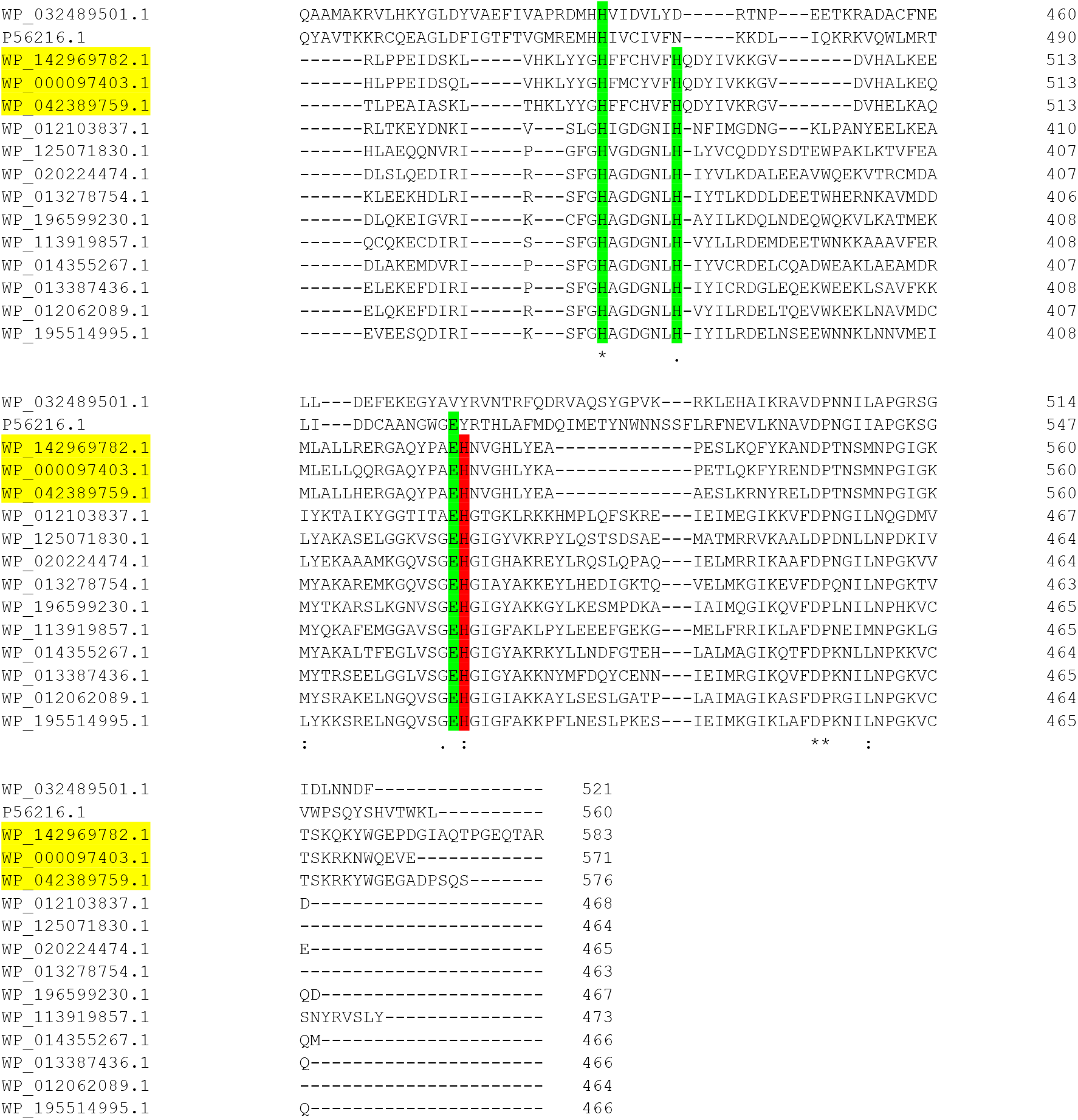
Sequence alignment of Ldh. WP_032489501.1, p-cresol methylhydroxylase of *Pseudomonas putida;* P56216.1, vanillyl-alcohol oxidase of *Penicillium simplicissimum;* WP_142969782.1, membrane-bound D-lactate dehydrogenase of *Cronobacter sakazakii;* WP_000097403.1, *Escherichia coli;* WP_042389759.1, *Pseudescherichia vulneris;* WP_012062089.1 (highlighted in yellow); bifurcating lactate dehydrogenase of *Alkaliphilus metalliredigens*; WP_125071830.1, *Lactiplantibacillus garii*; WP_020224474.1, *Holdemania massiliensis*; WP_013278754.1, *Acetohalobium arabaticum*; WP_196599230.1, *Pectinatus frisingensis*; WP_113919857.1, *Alkalibaculum bacchi*; WP_014355267.1, *A. woodii;* WP_013387436.1, *Ilyobacter polytropus;* WP_012103837.1, *Clostridium kluyveri;* WP_195514995.1, *Paraclostridium bifermentans*. The postulated acid catalyst His423 is marked in red, the residues ligating the metal ion in green and Asp39 and Asp137 in cyan.

**Table 1 – Figure supplement 1A.**
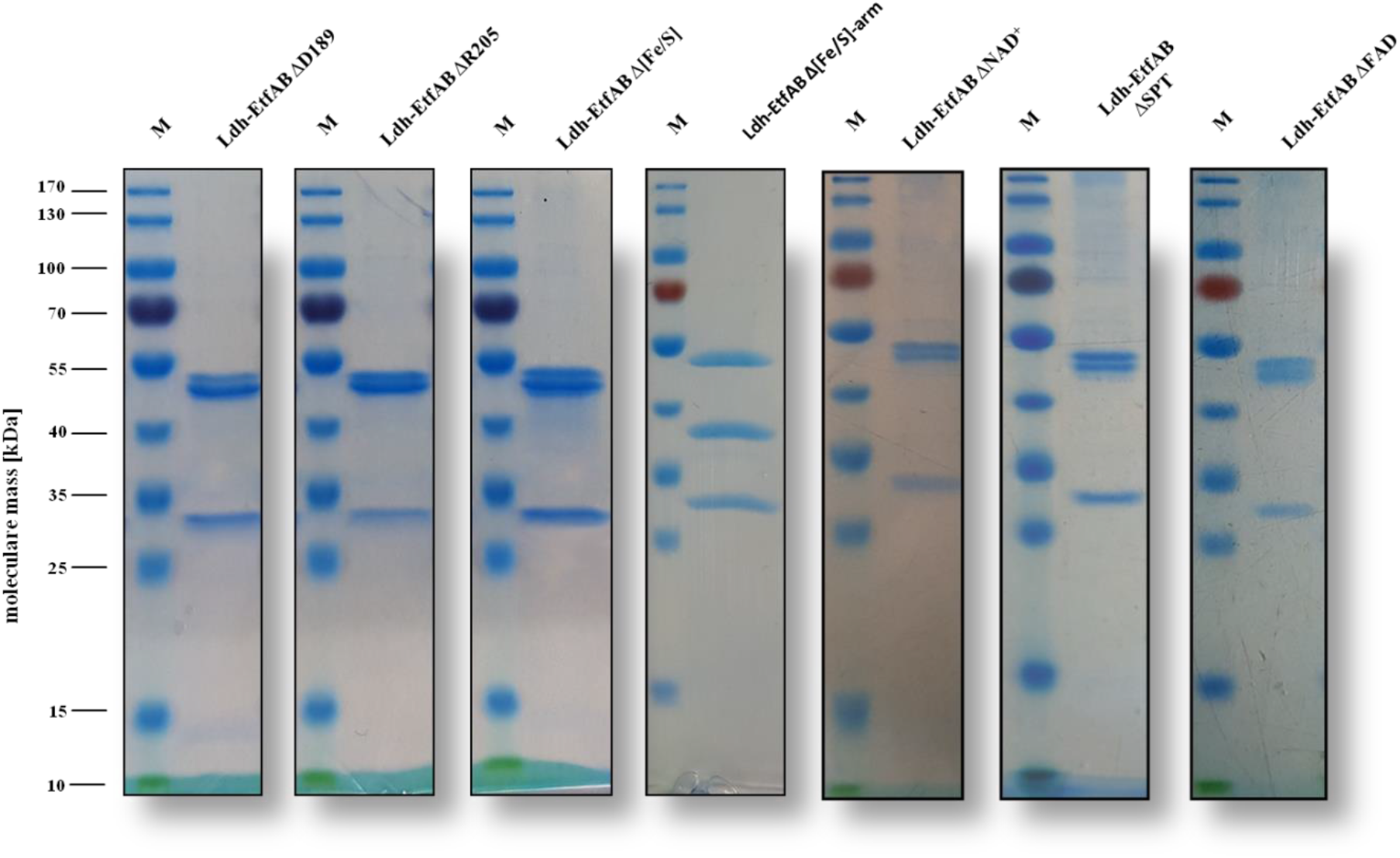
SDS-PAGE of the site-specifically changed Ldh-EtfAB variants. The Ldh-EtfAB variants were anaerobically purified by strep-tactin affinity and Superdex 200 size exclusion chromatography. 5 μg of the purified protein was separated by a 12% SDS-PAGE. Coomassie Brilliant Blue G250 was used for protein staining. M, Prestained PageRuler^™^ **Source data 1.** Source data for Figure supplement 1A

**Table 1 – Figure supplement 1B.**
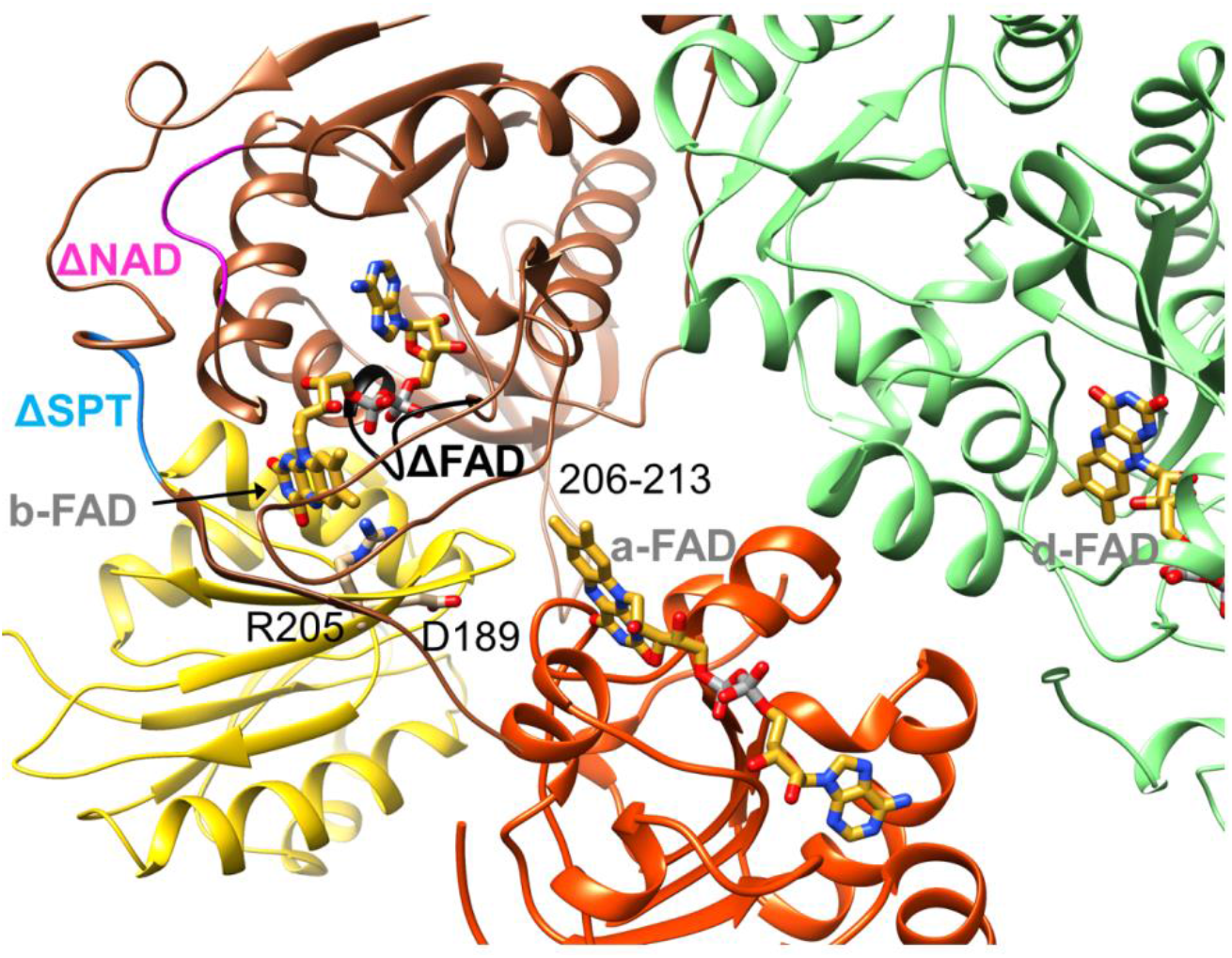
Location of the EtfA and EtfB mutations. The residues of the ΔFAD, ΔNAD and ΔSPT variants are highlighted in blacks, magenata and blue. The positions of R205 and D189 are shown as ball-and-sticks. The EtfAB shuttle is presented in B-state calculated by Alphafold2.

## Supplementary File 1

### A. Table of corresponding primers used

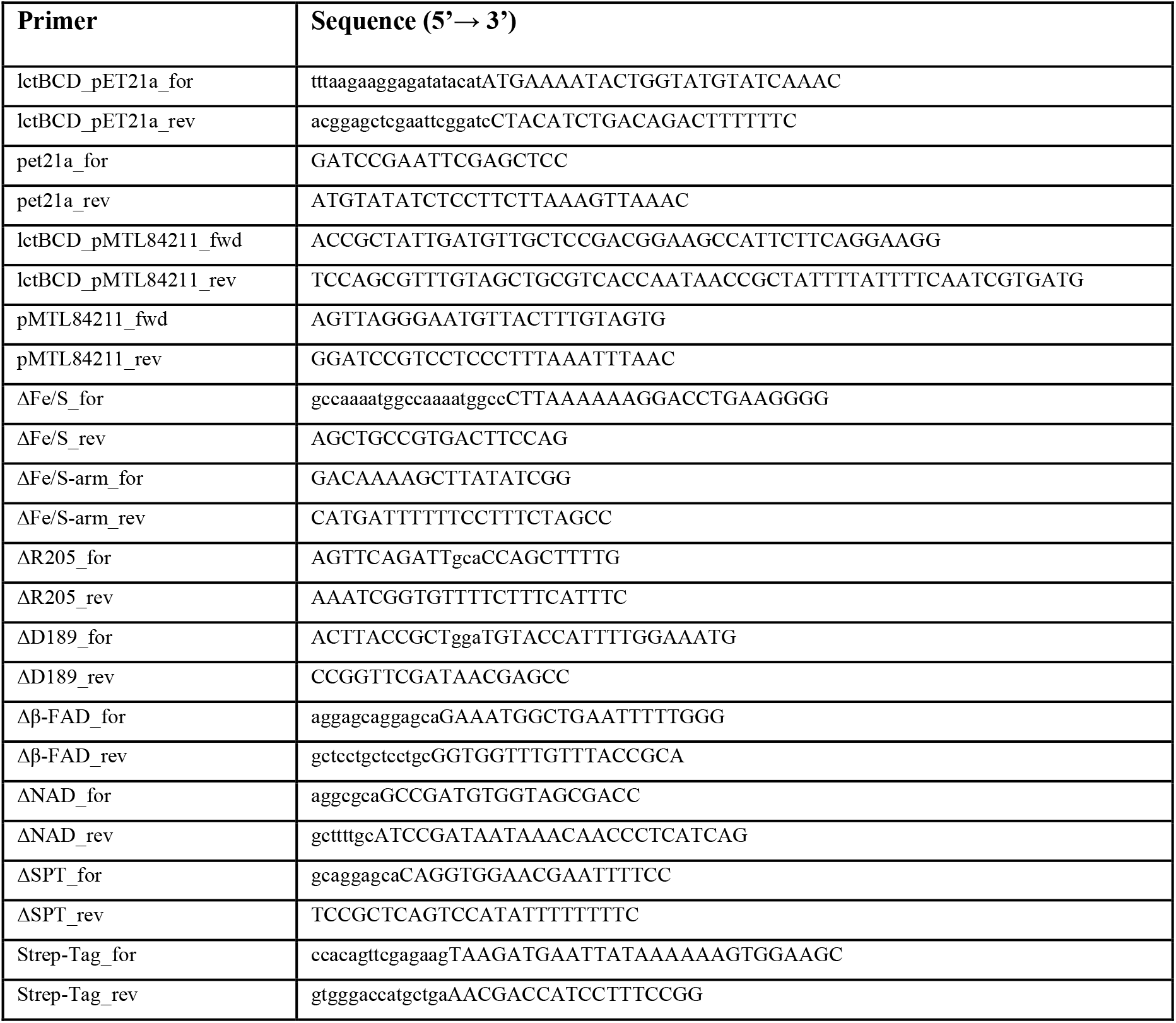

Plasmids generated with exchanges in EtfA are pET21a_lctBCD[ΔFe/S](C41A, C44A, C47A), pET21a_lctBCD[ΔFe/S-arm](Δ2A-65I), pMTL84211_lctBCD[ΔFe/S](C41A, C44A, C47A), pMTL84211_lctBCD[ΔFe/S-arm](Δ2A-65I), pET21a_lctBCD[ΔR205](R205A) and pET21a_lctBCD[ΔD189](D189A). Plasmids generated with exchanges in EtfB are pET21a_lctBCD[Δβ-FAD](D122A, D124A, T125G, Q127G, V128A, P130A), pET21a_lctBCD[ΔNAD^+^](R87A, F89A, G91A) and pET21a_lctBCD[ΔSPT](S223A, P224G, T225A).

### B. Cloning of *pET21a_lctBC-StrepD*

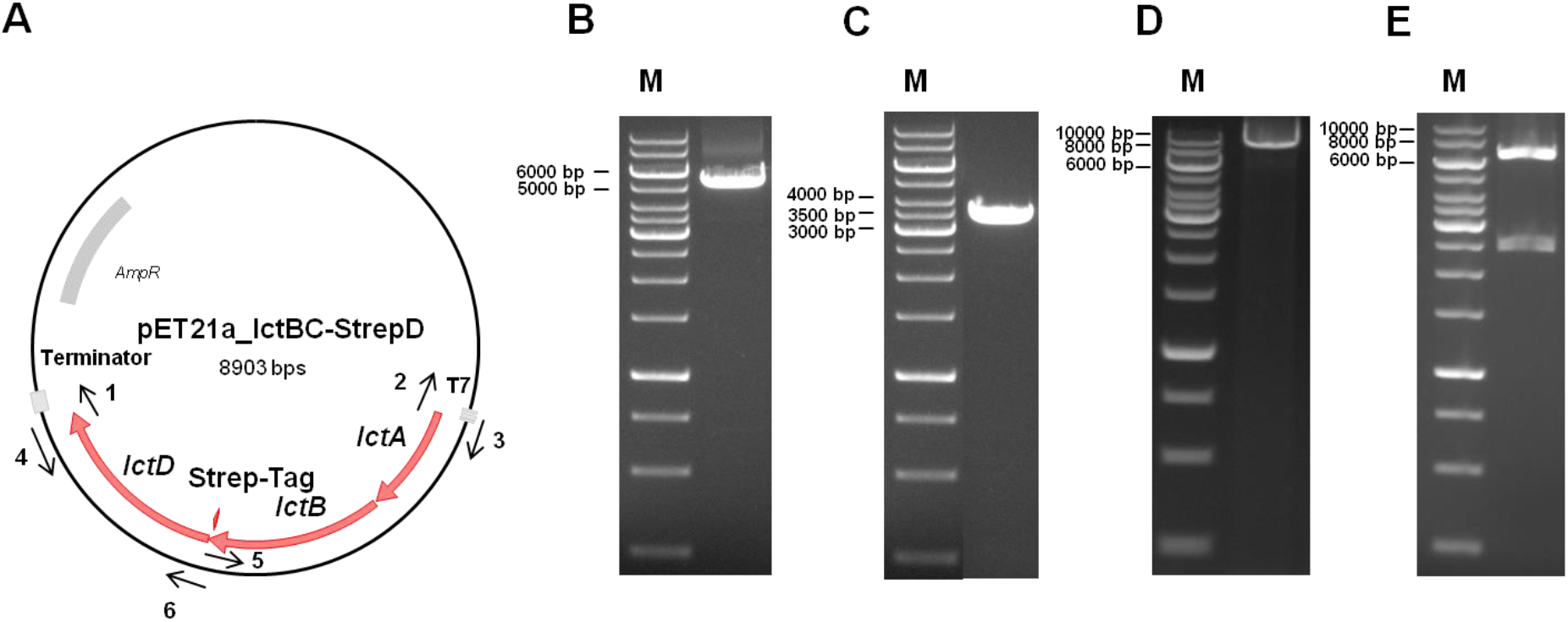

For the production of Ldh-EtfAB-Strep in *E. coli* the construct *pET21a_lctBC-StrepD* was cloned (A). Therefore, *pET21a* backbone, including a T7-promotor, was amplified using corresponding primers pET21a_for (1) and pET21a_rev (2) via PCR (size: 5406 bp) (B). *LctBCD* was amplified from genomic DNA of *A. woodii* via PCR, using lctBCD_pET21a_for (3) and lctBCD_pET21a_rev (4) primers (size: 3537 bp) (C). Amplified *lctBCD* and *pET21a* backbone were fused via Gibson Assembly and transformed in *E. coli* HB101. Afterwards, plasmids where isolated and a sequence encoding for a Strep-tag was introduced at the 3’-end of the gene *IctC* by using corresponding primers Strep-Tag_for (5) and Strep-Tag_rev (6) (size: 8903 bp) (D). The resulting *pET21a_lctBC-StrepD* was digested with *HindIII* (E). The resulting sizes were 6393 bp and 2510 bp. M, Gene Ruler 1 kb DNA ladder.

**Source data 1.** Source data for Supplementary Figure 1B

## References

Adams, P. D., Afonine, P. V., Bunkoczi, G., Chen, V. B., Davis, I. W., Echols, N., Headd, J. J., Hung, L. W., Kapral, G. J., Grosse-Kunstleve, R. W., McCoy, A. J., Moriarty, N. W., Oeffner, R., Read, R. J., Richardson, D. C., Richardson, J. S., Terwilliger, T. C., & Zwart, P. H. (2010). PHENIX: a comprehensive Python-based system for macromolecular structure solution [Research Support, N.I.H., Extramural Research Support, Non-U.S. Gov’t Research Support, U.S. Gov’t, Non-P.H.S.]. Acta Crystallogr D Biol Crystallogr, 66(Pt 2), 213–221. https://doi.org/10.1107/S0907444909052925

Baymann, F., Schoepp-Cothenet, B., Duval, S., Guiral, M., Brugna, M., Baffert, C., Russell, M. J., & Nitschke, W. (2018). On the natural history of flavin-based electron bifurcation. Front Microbiol, 9, 1357. https://doi.org/10.3389/fmicb.2018.01357

Benson, T. E., Walsh, C. T., & Hogle, J. M. (1997). X-ray crystal structures of the S229A mutant and wild-type MurB in the presence of the substrate enolpyruvyl-UDP-N-acetylglucosamine at 1.8-A resolution. Biochemistry, 36(4), 806–811. https://doi.org/10.1021/bi962221g

Bertsch, J., Parthasarathy, A., Buckel, W., & Müller, V. (2013). An electron-bifurcating caffeyl-CoA reductase [Research Support, Non–U.S. Gov’t]. J. Biol. Chem., 288(16), 11304–11311. https://doi.org/10.1074/jbc.M112.444919

Bock, A.-K., A., P.-K., & Schönheit, P. (1994). Pyruvate - a novel substrate for growth and methane formation in *Methanosarcina barkeri*. Archives of Microbiology, 161(1), 33–46.

Bradford, M. M. (1976). A rapid and sensitive method for the quantification of microgram quantities of protein utilizing the principle of proteine-dye-binding. Anal. Biochem., 72, 248–254.

Buckel, W., & Thauer, R. K. (2013). Energy conservation via electron bifurcating ferredoxin reduction and proton/Na(+) translocating ferredoxin oxidation [Research Support, Non–U.S. Gov’t Review]. Biochim Biophys Acta, 1827(2), 94–113. https://doi.org/10.1016/j.bbabio.2012.07.002

Buckel, W., & Thauer, R. K. (2018a). Flavin-based electron bifurcation, a new mechanism of biological energy coupling. Chem Rev, 118(7), 3862–3886. https://doi.org/10.1021/acs.chemrev.7b00707

Buckel, W., & Thauer, R. K. (2018b). Flavin-based electron bifurcation, ferredoxin, flavodoxin, and anaerobic respiration with protons (Ech) or NAD(+) (Rnf) as electron acceptors: a historical review. Front Microbiol, 9, 401. https://doi.org/10.3389/fmicb.2018.00401

Chowdhury, N. P., Mowafy, A. M., Demmer, J. K., Upadhyay, V., Koelzer, S., Jayamani, E., Kahnt, J., Hornung, M., Demmer, U., Ermler, U., & Buckel, W. (2014). Studies on the mechanism of electron bifurcation catalyzed by electron transferring flavoprotein (Etf) and butyryl-CoA dehydrogenase (Bcd) of *Acidaminococcus fermentans* [Research Support, Non–U.S. Gov’t]. J Biol Chem, 289(8), 5145–5157. https://doi.org/10.1074/jbc.M113.521013

Crofts, A. R., Hong, S., Wilson, C., Burton, R., Victoria, D., Harrison, C., & Schulten, K. (2013). The mechanism of ubihydroquinone oxidation at the Qo-site of the cytochrome *bc1* complex. Biochim Biophys Acta, 1827(11-12, 1362–1377. https://doi.org/10.1016/j.bbabio.2013.01.009

Cunane, L. M., Chen, Z. W., Shamala, N., Mathews, F. S., Cronin, C. N., & McIntire, W. S. (2000). Structures of the flavocytochrome p-cresol methylhydroxylase and its enzyme-substrate complex: gated substrate entry and proton relays support the proposed catalytic mechanism. J Mol Biol, 295(2), 357–374. https://doi.org/10.1006/jmbi.1999.3290

Demmer, J. K., Bertsch, J., Oppinger, C., Wohlers, H., Kayastha, K., Demmer, U., Ermler, U., & Müller, V. (2018). Molecular basis of the flavin-based electron-bifurcating caffeyl-CoA reductase reaction. FEBS Lett, 592(3), 332–342. https://doi.org/10.1002/1873-3468.12971

Demmer, J. K., Bertsch, J., Öppinger, C., Wohlers, H., Kayastha, K., Demmer, U., Ermler, U., & Müller, V. (2018). Molecular basis of the flavin-based electron-bifurcating caffeyl-CoA reductase reaction. FEBSLett. https://doi.org/InPress

Demmer, J. K., Huang, H., Wang, S., Demmer, U., Thauer, R. K., & Ermler, U. (2015). Insights into flavin-based electron bifurcation via the NADH-dependent reduced ferredoxin:NADP oxidoreductase structure [Research Support, Non–U.S. Gov’t]. J Biol Chem, 290(36), 21985–21995. https://doi.org/10.1074/jbc.M115.656520

Demmer, J. K., Pal Chowdhury, N., Selmer, T., Ermler, U., & Buckel, W. (2017a). The semiquinone swing in the bifurcating electron transferring flavoprotein/butyryl-CoA dehydrogenase complex from *Clostridium difficile*. Nat. Commun., 8, 1577. https://doi.org/10.1038/s41467-017-01746-3

Demmer, J. K., Pal Chowdhury, N., Selmer, T., Ermler, U., & Buckel, W. (2017b). The semiquinone swing in the bifurcating electron transferring flavoprotein/butyryl-CoA dehydrogenase complex from *Clostridium difficile*. Nat Commun, 8(1), 1577. https://doi.org/10.1038/s41467-017-01746-3

Dönig, J., & Müller, V. (2018). Alanine, a novel growth substrate for the acetogenic bacterium *Acetobacterium woodii*. Appl Environ Microbiol, 84(23). https://doi.org/10.1128/AEM.02023-18

Duan, H. D., Khan, S. A., & Miller, A. F. (2021). Photogeneration and reactivity of flavin anionic semiquinone in a bifurcating electron transfer flavoprotein. Biochim Biophys Acta Bioenerg, 1862(7), 148415. https://doi.org/10.1016/j.bbabio.2021.148415

Dym, O., Pratt, E. A., Ho, C., & Eisenberg, D. (2000). The crystal structure of D-lactate dehydrogenase, a peripheral membrane respiratory enzyme. Proc Natl Acad Sci U S A, 97(17), 9413–9418. https://doi.org/10.1073/pnas.97.17.9413

Emsley, P., & Cowtan, K. (2004). Coot: model-building tools for molecular graphics [Research Support, Non–U.S. Gov’t]. Acta Crystallogr D Biol Crystallogr, 60(Pt 12 Pt 1), 2126–2132. https://doi.org/10.1107/S0907444904019158

Feng, X., Schut, G. J., Lipscomb, G. L., Li, H., & Adams, M. W. W. (2021). Cryoelectron microscopy structure and mechanism of the membrane-associated electron-bifurcating flavoprotein Fix/EtfABCX. Proc Natl Acad Sci U S A, 118(2). https://doi.org/10.1073/pnas.2016978118

Fish, W. W. (1988). Rapid colorimetric micromethod for the quantitation of complexed iron in biological samples. Methods Enzymol., 158, 357–364. http://www.ncbi.nlm.nih.gov/entrez/query.fcgi?cmd=Retrieve&db=PubMed&dopt=Citation&list_uids=3374387

Fraaije, M. W., Van Berkel, W. J., Benen, J. A., Visser, J., & Mattevi, A. (1998). A novel oxidoreductase family sharing a conserved FAD-binding domain. Trends Biochem Sci, 23(6), 206–207. https://doi.org/10.1016/s0968-0004(98)01210-9

Garcia Costas, A. M., Poudel, S., Miller, A. F., Schut, G. J., Ledbetter, R. N., Fixen, K. R., Seefeldt, L. C., Adams, M. W. W., Harwood, C. S., Boyd, E. S., & Peters, J. W. (2017). Defining electron bifurcation in the electron-transferring flavoprotein family. J Bacteriol, 199(21). https://doi.org/10.1128/JB.00440-17

Jumper, J., Evans, R., Pritzel, A., Green, T., Figurnov, M., Ronneberger, O., Tunyasuvunakool, K., Bates, R., Zidek, A., Potapenko, A., Bridgland, A., Meyer, C., Kohl, S. A. A., Ballard, A. J., Cowie, A., Romera-Paredes, B., Nikolov, S., Jain, R., Adler, J.,… Hassabis, D. (2021). Highly accurate protein structure prediction with AlphaFold. Nature, 596(7873), 583–589. https://doi.org/10.1038/s41586-021-03819-2

Kandler, O. (1983). Carbohydrate metabolism in lactic acid bacteria. Antonie Van Leeuwenhoek, 49(3), 209–224. https://doi.org/10.1007/BF00399499

Katsyv, A., Jain, S., Basen, M., & Muller, V. (2021). Electron carriers involved in autotrophic and heterotrophic acetogenesis in the thermophilic bacterium Thermoanaerobacter kivui. Extremophiles. https://doi.org/10.1007/s00792-021-01247-8

Kayastha, K., Vitt, S., Buckel, W., & Ermler, U. (2021). Flavins in the electron bifurcation process. Arch Biochem Biophys, 701, 108796. https://doi.org/10.1016/i.abb.2021.108796

Laemmli, U. K. (1970). Cleavage of structural proteins during the assembly of the head of bacteriophage T4. Nature, 227(5259), 680–685. https://doi.org/10.1038/227680a0

Langer, G., Cohen, S. X., Lamzin, V. S., & Perrakis, A. (2008). Automated macromolecular model building for X-ray crystallography using ARP/wARP version 7. Nat Protoc, 3(7), 1171–1179. https://doi.org/10.1038/nprot.2008.91

Leys, D., Basran, J., Talfournier, F., Sutcliffe, M. J., & Scrutton, N. S. (2003). Extensive conformational sampling in a ternary electron transfer complex [Research Support, Non–U.S. Gov’t]. Nat Struct Biol, 10(3), 219–225. https://doi.org/10.1038/nsb894

Lubner, C. E., Jennings, D. P., Mulder, D. W., Schut, G. J., Zadvornyy, O. A., Hoben, J. P., Tokmina-Lukaszewska, M., Berry, L., Nguyen, D. M., Lipscomb, G. L., Bothner, B., Jones, A. K., Miller, A. F., King, P. W., Adams, M. W. W., & Peters, J. W. (2017). Mechanistic insights into energy conservation by flavin-based electron bifurcation. Nat Chem Biol, 13(6), 655–659. https://doi.org/10.1038/nchembio.2348

Mattevi, A., Fraaije, M. W., Mozzarelli, A., Olivi, L., Coda, A., & van Berkel, W. J. (1997). Crystal structures and inhibitor binding in the octameric flavoenzyme vanillyl-alcohol oxidase: the shape of the active-site cavity controls substrate specificity. Structure, 5(7), 907–920. https://doi.org/10.1016/s0969-2126(97)00245-1

Müller, V. (2008). Bacterial fermentation. In: John Wiley & Sons, Ltd: Chichester.

Müller, V., Chowdhury, N. P., & Basen, M. (2018). Electron bifurcation: A long-hidden energy-coupling mechanism. Annu Rev Microbiol, 72, 331–353. https://doi.org/10.1146/annurev-micro-090816-093440

Nitschke, W., & Russell, M. J. (2012). Redox bifurcations: mechanisms and importance to life now, and at its origin: a widespread means of energy conversion in biology unfolds. Bioessays, 34(2), 106–109. https://doi.org/10.1002/bies.201100134

Peters, J. W., Miller, A. F., Jones, A. K., King, P. W., & Adams, M. W. (2016). Electron bifurcation [Review]. Curr Opin Chem Biol, 31, 146–152. https://doi.org/10.1016/j.cbpa.2016.03.007

Pettersen, E. F., Goddard, T. D., Huang, C. C., Couch, G. S., Greenblatt, D. M., Meng, E. C., & Ferrin, T. E. (2004). UCSF Chimera--a visualization system for exploratory research and analysis. J Comput Chem, 25(13), 1605–1612. https://doi.org/10.1002/jcc.20084

Punjani, A., Rubinstein, J. L., Fleet, D. J., & Brubaker, M. A. (2017). cryoSPARC: algorithms for rapid unsupervised cryo-EM structure determination. Nat Methods, 14(3), 290–296. https://doi.org/10.1038/nmeth.4169

Roberts, D. L., Frerman, F. E., & Kim, J. J. (1996). Three-dimensional structure of human electron transfer flavoprotein to 2.1-A resolution [Research Support, U.S. Gov’t, P.H.S.]. Proc Natl Acad Sci USA, 93(25), 14355–14360. http://www.ncbi.nlm.nih.gov/pubmed/8962055

Scheres, S. H. (2012). RELION: implementation of a Bayesian approach to cryo-EM structure determination. J Struct Biol, 180(3), 519–530. https://doi.org/10.1016/i.isb.2012.09.006

Schönheit, P., Wäscher, C., & Thauer, R. K. (1978). A rapid procedure for the purification of ferredoxin from Clostridia using polyethylenimine. FEBS Lett., 89, 219–222.

Schuchmann, K., & Müller, V. (2012). A bacterial electron bifurcating hydrogenase. J. Biol. Chem., 287, 31165–31171. http://www.ncbi.nlm.nih.gov/entrez/query.fcgi?cmd=Retrieve&db=PubMed&dopt=Citation&list_uids=22810230

Schut, G. J., & Adams, M. W. (2009). The iron-hydrogenase of *Thermotoga maritima* utilizes ferredoxin and NADH synergistically: a new perspective on anaerobic hydrogen production. J. Bacteriol., 191(1), 4451–4457. https://doi.org/JB.01582-08[pii]10.1128/JB.01582-08

Schut, G. J., Mohamed-Raseek, N., Tokmina-Lukaszewska, M., Mulder, D. W., Nguyen, D. M. N., Lipscomb, G. L., Hoben, J. P., Patterson, A., Lubner, C. E., King, P. W., Peters, J. W., Bothner, B., Miller, A. F., & Adams, M. W. W. (2019a). The catalytic mechanism of electron-bifurcating electron transfer flavoproteins (ETFs) involves an intermediary complex with NAD(). J Biol Chem, 294(9), 3271–3283. https://doi.org/10.1074/jbc.RA118.005653

Schut, G. J., Mohamed-Raseek, N., Tokmina-Lukaszewska, M., Mulder, D. W., Nguyen, D. M. N., Lipscomb, G. L., Hoben, J. P., Patterson, A., Lubner, C. E., King, P. W., Peters, J. W., Bothner, B., Miller, A. F., & Adams, M. W. W. (2019b). The catalytic mechanism of electron-bifurcating electron transfer flavoproteins (ETFs) involves an intermediary complex with NAD^+^. J Biol Chesm, 294(9), 3271–3283. https://doi.org/10.1074/jbc.RA118.005653

Sucharitakul, J., Buttranon, S., Wongnate, T., Chowdhury, N. P., Prongjit, M., Buckel, W., & Chaiyen, P. (2020). Modulations of the reduction potentials of flavin-based electron bifurcation complexes and semiquinone stabilities are key to control directional electron flow. FEBS J. https://doi.org/10.1111/febs.15343

Toogood, H. S., Leys, D., & Scrutton, N. S. (2007). Dynamics driving function: new insights from electron transferring flavoproteins and partner complexes [Research Support, Non–U.S. Gov’t Review]. FEBS J, 274(21), 5481–5504. https://doi.org/10.1111/j.1742-4658.2007.06107.x

Vita, N., Valette, O., Brasseur, G., Lignon, S., Denis, Y., Ansaldi, M., Dolla, A., & Pieulle, L. (2015). The primary pathway for lactate oxidation in *Desulfovibrio vulgaris*. Front Microbiol, 6, 606. https://doi.org/10.3389/fmicb.2015.00606

Wagner, T., Koch, J., Ermler, U., & Shima, S. (2017). Methanogenic heterodisulfide reductase (HdrABC-MvhAGD) uses two noncubane [4Fe-4S] clusters for reduction. Science, 357(6352), 699–703. https://doi.org/10.1126/science.aan0425

Wagner, T., Merino, F., Stabrin, M., Moriya, T., Antoni, C., Apelbaum, A., Hagel, P., Sitsel, O., Raisch, T., Prumbaum, D., Quentin, D., Roderer, D., Tacke, S., Siebolds, B., Schubert, E., Shaikh, T. R., Lill, P., Gatsogiannis, C., & Raunser, S. (2019). SPHIRE-crYOLO is a fast and accurate fully automated particle picker for cryo-EM. Commun Biol, 2, 218. https://doi.org/10.1038/s42003-019-0437-z

Wang, S., Huang, H., Kahnt, J., & Thauer, R. K. (2013). A reversible electron-bifurcating ferredoxin- and NAD-dependent [FeFe]-hydrogenase (HydABC) in *Moorella thermoacetica* [Research Support, Non–U.S. Gov’t]. J. Bacteriol., 195(6), 1267–1275. https://doi.org/10.1128/JB.02158-12

Watanabe, T., Pfeil-Gardiner, O., Kahnt, J., Koch, J., Shima, S., & Murphy, B. J. (2021). Three-megadalton complex of methanogenic electron-bifurcating and CO2-fixing enzymes. Science, 373(6559), 1151–1156. https://doi.org/10.1126/science.abg5550

Weghoff, M. C., Bertsch, J., & Müller, V. (2015). A novel mode of lactate metabolism in strictly anaerobic bacteria. Environ. Microbiol., 17, 670–677.http://www.ncbi.nlm.nih.gov/pubmed/24762045

Williams, C. J., Headd, J. J., Moriarty, N. W., Prisant, M. G., Videau, L. L., Deis, L. N., Verma, V., Keedy, D. A., Hintze, B. J., Chen, V. B., Jain, S., Lewis, S. M., Arendall, W. B., 3rd, Snoeyink, J., Adams, P. D., Lovell, S. C., Richardson, J. S., & Richardson, D. C. (2018). MolProbity: More and better reference data for improved all-atom structure validation. Protein Sci, 27(1), 293–315. https://doi.org/10.1002/pro.3330

Wittig, I., Carrozzo, R., Santorelli, F. M., & Schägger, H. (2007). Functional assays in high-resolution clear native gels to quantify mitochondrial complexes in human biopsies and cell lines. Electrophoresis, 28, 3811–3820.

Zhang, K. (2016). Gctf: Real-time CTF determination and correction. J Struct Biol, 193(1), 112. https://doi.org/10.1016/i.jsb.2015.11.003

Zheng, S. Q., Palovcak, E., Armache, J. P., Verba, K. A., Cheng, Y., & Agard, D. A. (2017). MotionCor2: anisotropic correction of beam-induced motion for improved cryoelectron microscopy. Nat Methods, 14(4), 331–332. https://doi.org/10.1038/nmeth.4193

Zivanov, J., Nakane, T., Forsberg, B. O., Kimanius, D., Hagen, W. J., Lindahl, E., & Scheres, S. H. (2018). New tools for automated high-resolution cryo-EM structure determination in RELION-3. Elife, 7. https://doi.org/10.7554/eLife.42166

